# Pain and itch processing by subpopulations of molecularly diverse spinal and trigeminal projection neurons

**DOI:** 10.1101/2020.06.17.156091

**Authors:** R. Wercberger, J.M. Braz, J.A. Weinrich, A.I. Basbaum

## Abstract

A remarkable molecular and functional heterogeneity of the primary sensory neurons and dorsal horn interneurons transmits pain- and or itch-relevant information, but the molecular signature of the projection neurons that convey the messages to the brain is unclear. Here, using retro-TRAP (translating ribosome affinity purification) and RNA-seq we reveal extensive molecular diversity of spino- and trigeminoparabrachial projection neurons, which to date are almost exclusively defined by their expression of the neurokinin 1 receptor (NK1R). Among the many genes identified, we highlight distinct subsets of *Cck*+, *Nptx2*+, *Nmb*+, and *Crh*+ expressing projection neurons. By combining *in situ* hybridization of retrogradely labeled neurons with Fos-based assays we also demonstrate significant functional heterogeneity, including both convergence and segregation of pain- and itch-provoking inputs onto molecularly diverse subsets of NK1R- and non-NK1R-expressing projection neurons. The current study provides the first comprehensive investigation into the molecular profiles and functional properties of projection neuron subtypes.

## INTRODUCTION

Recent studies described a plethora of neurochemically-distinct primary afferent and spinal cord interneuron populations that are tuned to discrete pain and itch stimulus modalities (Braz et al., 2014; Koch et al., 2018; LaMotte et al., 2014; Peirs and Seal, 2016; Todd, 2010). Importantly, however, the generation of modality specific percepts does not arise from the brain’s analysis of the activity of interneurons, but rather from activity of different projection neuron populations, which must be interpreted at supraspinal loci. It is, therefore, critical to address the extent to which the primary afferent and interneuron heterogeneity and specificity extends to the projection neurons in the spinal cord and trigeminal nucleus caudalis (TNC) (Wercberger and Basbaum, 2019).

Anatomical studies identified three morphological classes of lamina I projection neurons, and there is some, albeit controversial, evidence for the correlation of morphology with the electrophysiological profile, ascending projection and receptor expression of these cells (for review see Todd, 2010). There is certainly considerable evidence for functional heterogeneity. For example, many lamina I projection neurons respond only to noxious mechanical and/or thermal stimulation (Bester et al., 2000; Christensen and Perl, 1970; Craig et al., 2001; Hachisuka et al., 2020, 2016; Han et al., 1998). Another population, in primates (Craig et al., 2001; Dostrovsky and Craig, 1996) and cats (Han et al., 1998; Light et al., 1993), responds only to innocuous cooling. Lastly, there is a significant population of wide dynamic range (WDR) neurons that respond to both innocuous and noxious stimuli in the deep dorsal horn (Bester et al., 2000; Craig and Kniffki, 1985; Hylden et al., 1986; Wall, 1967; Willis Jr., 1985). Subsets of the WDR and nociceptive-specific neurons are pruriceptive (Carstens, 1997; Davidson et al., 2012, 2007; Jansen and Giesler, 2015; Moser and Giesler, 2014; Simone et al., 2004), with separate populations, in primates, responding to the pruritogens, histamine and cowage (Davidson et al., 2012, 2007).

From a molecular perspective, however, the projection neurons are often regarded as relatively homogeneous. With few exceptions, the neurokinin 1 receptor (NK1R) defines projection neurons in lamina I and the lateral spinal nucleus (LSN) (Carstens et al., 2010; Mantyh et al., 1997). On the other hand, there is evidence for subsets of NK1R neurons, based on projection target (Huang et al., 2019) or molecular make-up (Blomqvist and Mackerlova, 1995; Cameron et al., 2015; Gamboa-Esteves et al., 2004; Huang et al., 2019). Most recently, using unbiased single cell transcriptomics, Häring et al. (2018) identified an excitatory neuron cluster (Glut15) that includes NK1R-expressing spinoparabrachial neurons. This cluster included *Lypd1*, a forebrain protein implicated in anxiety disorders (Tekinay et al., 2009). However, as *Lypd1* labels ∼95% of spinoparabrachial neurons (Häring et al., 2018), it likely does not define a functionally distinct subset.

The approach that we took involved a sequential series of filtering steps. First, we performed projection neuron-centric RNA-sequencing. Using retro-TRAP (Translating Ribosome Affinity Purification), we purified lateral parabrachial (LPb)-projecting neurons from the spinal cord and TNC, and generated RNA-seq datasets of candidate projection neuron marker genes. As many of these genes are also expressed by spinal cord and TNC interneurons, in the next filtering step we verified the projection neuron hits by combining retrograde tracing and multiplexed in situ hybridization. The latter studies were qualitative in nature, but they identified genes that establish molecular heterogeneity of projection neurons. Lastly, we performed functional studies using TRAP2 mice (Targeted Recombination in Active Populations; Allen et al., 2017) and Fos in response to pain- and itch-provoking stimuli and demonstrated functional heterogeneity in the molecular diverse projection neuron population. Taken together, we report both algogen and pruritogen convergence as well as preliminary evidence for labeled-line properties of molecularly diverse subsets of NK1R- and non-NK1R-expressing projection neurons.

## RESULTS

### Selective purification and profiling of projection neuron RNA

To purify and sequence RNA specifically from projection neurons we injected a replication deficient, retrograde herpes simplex virus (HSV)-based viral vector encoding a green fluorescent protein (GFP)-tagged large ribosomal subunit protein L10 (HSV-GFPL10) into the LPb of wildtype mice, which induced expression of GFP-L10 in spino- and trigeminoparabrachial projection neurons (**Fig. 1A**). In a parallel study, to selectively target LPb-projecting neurons that express the NK1R, we injected an HSV-based viral vector encoding a Cre-recombinase dependent HA-tagged L10 (HSV-flex-HAL10) into the LPb of NK1R-cre mice. Data obtained from animals injected with HSV-GFPL10 or HSV-flex-HAL10 are hereafter referred to as the “PN” or “NK” dataset, respectively. To increase the total number of labeled projection neurons, in the following studies we combined spinal cord and trigeminal tissue. Two weeks after the viral injections, we recorded GFP- and HA-tagged ribosomes in projection neurons throughout the spinal cord and TNC in wildtype (**Fig. 1B**) and in NK1R-Cre (**Fig. 1C**) mice, respectively. Confirmation of specific immunolabeling of projection neurons was followed by the immunoprecipitation of GFP- or HA-tagged ribosomes and associated mRNA from spinal cord and trigeminal tissue.

**Figure 1:**
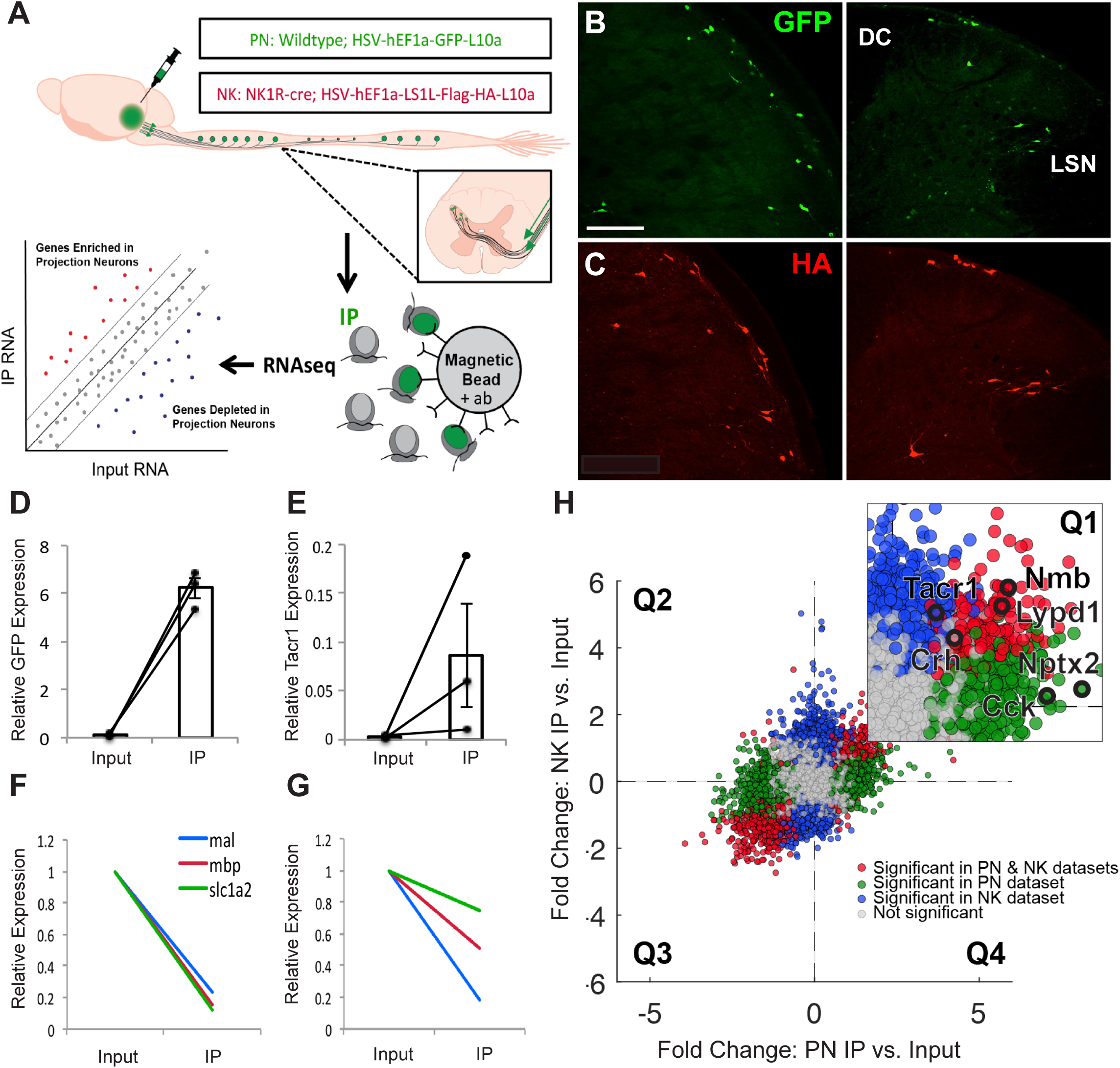
Selective purification and profiling of projection neurons reveals candidates for projection neuron marker genes. **(A)** Experimental design **(B**,**C)** Representative images of GFP (**B**) and HA (**C**) immunofluorescence in nucleus caudalis (left) and spinal cord (right) from wildtype and NK1R-Cre mice, respectively, illustrate projection neurons GFP or HA-tagged ribosomal protein. Scale bar 100μm. **(D**,**E)** qPCR results showing enrichment of *Gfp* **(D)** and *Tacr1* **(E)** in IP relative to Input samples, in PN and NK experiments, respectively. Data are normalized to Rpl27 and represented as mean ± SEM. **(F**,**G)** qPCR shows depletion of glial genes in IP relative to Input samples in PN **(F)** and NK **(G)** experiments. Data are normalized to Rpl27 and Input relative expression and represented as mean ± SEM. **(H)** RNA sequencing shows differential expression data of IP relative to Input Fold Change for PN experiments versus NK experiments. Quadrant 1 (Q1) contains genes enriched in both datasets, Q2 contains genes depleted in both, Q3 and Q4 contain genes differentially altered in PN versus NK datasets. Inset shows enlarged Q1 with genes of interest highlighted in black. Genes significantly enriched or depleted in both PN and NK datasets are highlighted in red. Genes significantly changed in PN, but not NK dataset are highlighted in green, while genes significantly changed in NK, but not PN, dataset are highlighted in blue. All significant differences P<.05.

Specificity of the immunoprecipitation (IP) was confirmed by Taqman qPCR. In all IP versus Input libraries we recorded enrichment of *Gfp* (∼64.6-fold; **Fig. 1D, Supplementary Fig. 1A**) and *Tacr1* (∼26.5-fold; **Fig. 1E, Supplementary Fig. 1B**). Depletion of glial markers in the PN libraries (*mbp*: ∼6.4-fold; *mal:* ∼3.9-fold; *slc1a2:* ∼7.0-fold) and in the NK libraries (*mbp:* ∼4.2-fold; *mal:* ∼5.4-fold; *slc1a2:* ∼1.2-fold) (**Fig. 1F,G** and **Supplementary Fig. 1C**) further confirmed specificity of the IPs.

### Candidate projection neuron marker genes

After qPCR confirmation that the IPs were specific for projection neurons, as a first filtering step, we performed bulk ribosomal RNA-sequencing on all IP (GFP or HA-tagged ribosomes) and input samples (dorsal spinal cord and TNC). To compare the PN and NK datasets obtained by RNA sequencing and differential expression analysis, (**Supplementary Figs. 1F,G**; **Supplementary Files 1,2**) we plotted the gene fold changes within each dataset against the changes in the other (**Fig. 1H**). Data points in quadrants one (Q1) and three (Q3) represent transcripts that are enriched or depleted, respectively, in projection neurons in both datasets; data points in quadrants two (Q2) and four (Q4) represent transcripts that are differentially enriched or depleted. As ∼90% of all lamina I spinoparabrachial neurons express the NK1R (Al-Khater and Todd, 2009; Spike et al., 2003), we expected both datasets to be largely overlapping and thus the majority of data points should lie in Q1 and Q3. This was indeed what we observed. Note, however, that the transcripts in Q4 provide a valuable (albeit limited) list of potential markers for the non-NK1R expressing projection neurons, for which there are currently no or only very limited marker genes identified (**Supplementary File 3).** Lastly, using qPCR we confirmed enrichment of the candidate marker-genes hits from the RNA-seq datasets (**Supplementary Figs. 1D,E**).

Because the first filter (RNA-sequencing and confirmation by qPCR) identified pan-neuronal as well as projection neuron markers, in a second filter step we performed fluorescent ISH to narrow down the list of hits to those more discretely expressed by projection neurons. As a result, in the remainder of this report, we focused on several genes that are enriched in both PN and NK RNA-seq and qPCR datasets (Q1, **Fig. 1H, Supplementary Figs. 1D-G**) and that have previously been implicated in pain and/or itch processing but not emphasized with respect to projection neuron neurochemistry.

*Cck* encodes cholecystokinin, a peptide expressed in dorsal horn neurons (Bras et al., 1999; Elde et al., 1990; Gutierrez-Mecinas et al., 2019; Häring et al., 2018; Hökfelt et al., 2001; Leah et al., 1988) and has been implicated as an anti-opioid (Faris et al., 1983; Watkins et al., 1985). To confirm our RNASeq and qPCR findings of *Cck* enrichment in projection neurons we retrogradely labeled LPb-projecting neurons with HSV-GFPL10 and performed double fluorescent ISH for *Gfp* and *Cck* in the TNC. Figure 2A illustrates *Cck*-expressing*/Gfp*-positive (i.e. projection neurons) in laminae II, III and IV, and interestingly, many fewer in lamina I (**Fig. 2A**). However, based on the high number of *Cck-*positive*/Gfp-*negative neurons, we conclude that the majority of the *Cck*-expressing cells are interneurons. The neuronal pentraxin 2 (*Nptx2*) gene encodes a secreted protein involved in excitatory synaptogenesis. *Nptx*2 has been implicated in various neuropsychiatric disorders (Chang et al., 2018) and pain processing (Miskimon et al., 2014). Here we recovered *Gfp*-expressing LPb–projecting neurons that co-express *Nptx2*, predominantly in laminae I and III/IV (**Fig. 2B**). As for *Cck*, however, the high number of *Nptx2*-positive/*Gfp-*negative neurons indicates that *Nptx2-*expresssing interneurons predominate. Neuromedin B (*Nmb*), a member of the bombesin-like family of peptides is robustly expressed in sensory neurons and in scattered dorsal horn neurons (Fleming et al., 2012) and has been implicated in both pain and itch processing (Fleming et al., 2012; Mishra et al., 2012; Su and Ko, 2011; Wan et al., 2017). We found that *Nmb* is expressed sparsely in the TNC, predominantly in superficial laminae and a subset of these were *Gfp*-labeled LPb-projecting neurons (**Fig. 2C**). Lastly, we characterized corticotropin-releasing hormone (*Crh*), which is a major contributor to the HPA axis-induced stress response and has been implicated in both peripheral and central pain processing (Hargreaves et al., 1989; Lariviere and Melzack, 2000). We recorded *Crh*/*Gfp*-labeled LPb-projecting neurons (**Fig. 2D**) in superficial TNC. Taken together, these results identify novel markers of LPb-projecting neurons. Interestingly, although Häring et al. (2018) reported that *Crh* is expressed in the Glut15 cluster, which includes spinoparabrachial neurons, this was not the case for *Cck, Nptx2* and *Nmb*.

**Figure 2:**
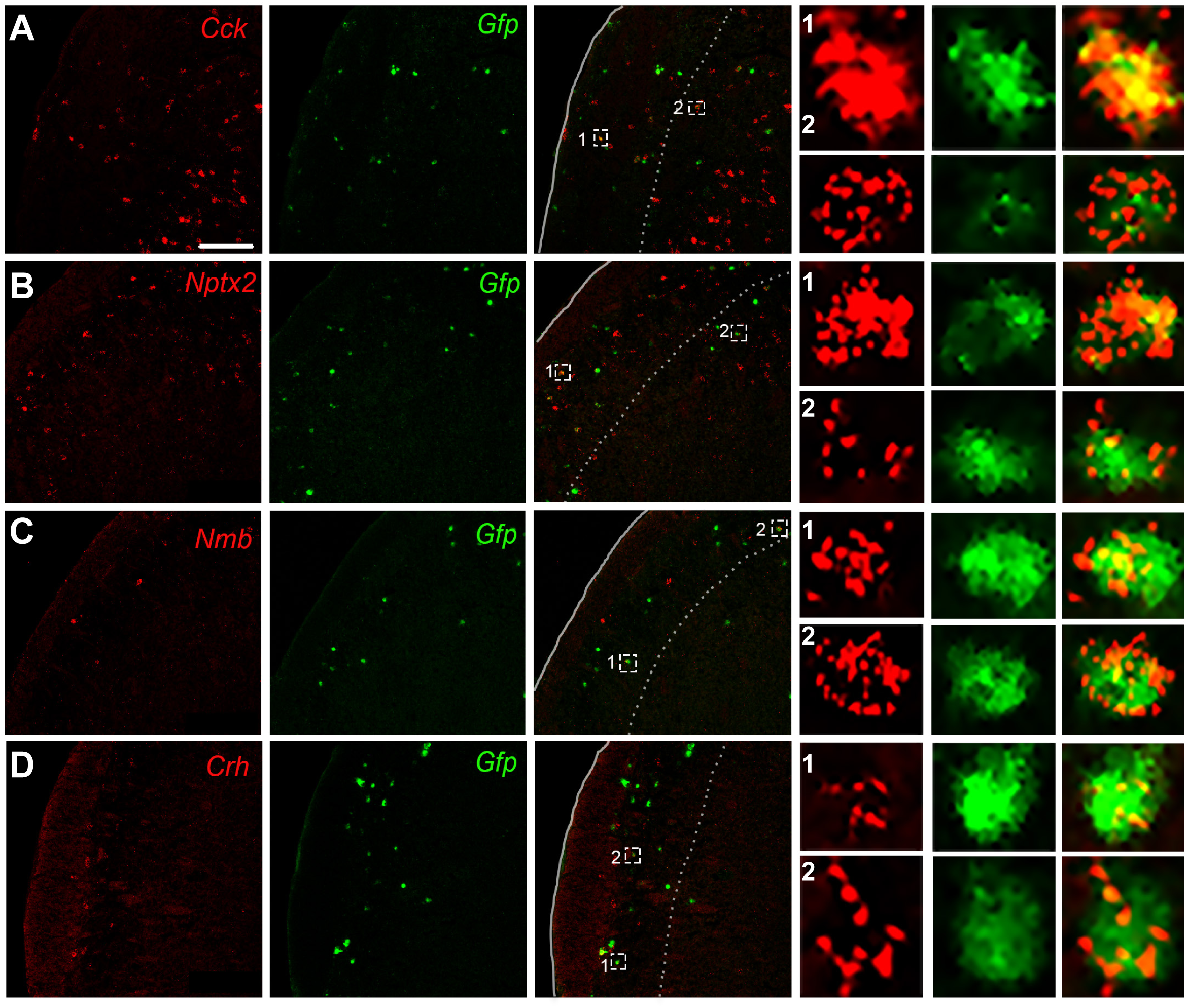
*In situ* hybridization confirms projection neuron marker genes identified by RNA-seq. Representative TNC sections illustrate co-labeling of *Cck* **(A)**, *Nptx2* **(B)**, *Nmb* **(C)**, and *Crh* **(D)** mRNA (red) with *Gfp*-tagged trigeminoparabrachial projection neurons (green). Insets show enlarged examples of individual cells positive for both the marker gene and *Gfp*. Scale bars, 100μm.

### Molecular heterogeneity of the NK1R-expressing dorsal horn neurons

To determine the extent to which these projection neuron markers constitute subsets of the NK1R-expressing projection neurons, or whether they define unique populations, we next performed double- and triple-fluorescent ISH for *Tacr1* and each of the enriched genes (**Fig. 3, Supplementary Fig. 2**). In fact, for each gene tested we recorded cells that co-expressed the enriched gene and *Tacr1* and others only expressed the gene or *Tacr1*. Interestingly, although we observed *Tacr1*-expressing neurons that co-expressed *Cck* in the deep dorsal horn, only rarely did we find *Cck* and *Tacr1* coexpressed in lamina I neurons (**Supplementary Fig. 2A**). In both superficial and deep dorsal horn, we observed subsets of *Tacr1*-expressing neurons that coexpress *Nptx2* (**Supplementary Fig. 2B**), *Nmb* (**Supplementary Fig. 2C**), and *Crh* (**Supplementary Fig. 2D**). In all cases, we observed many neurons that solely expressed *Tacr1*, or that were positive for the marker gene, and not *Tacr1* (**Supplementary Fig. 2A-D**).

**Figure 3:**
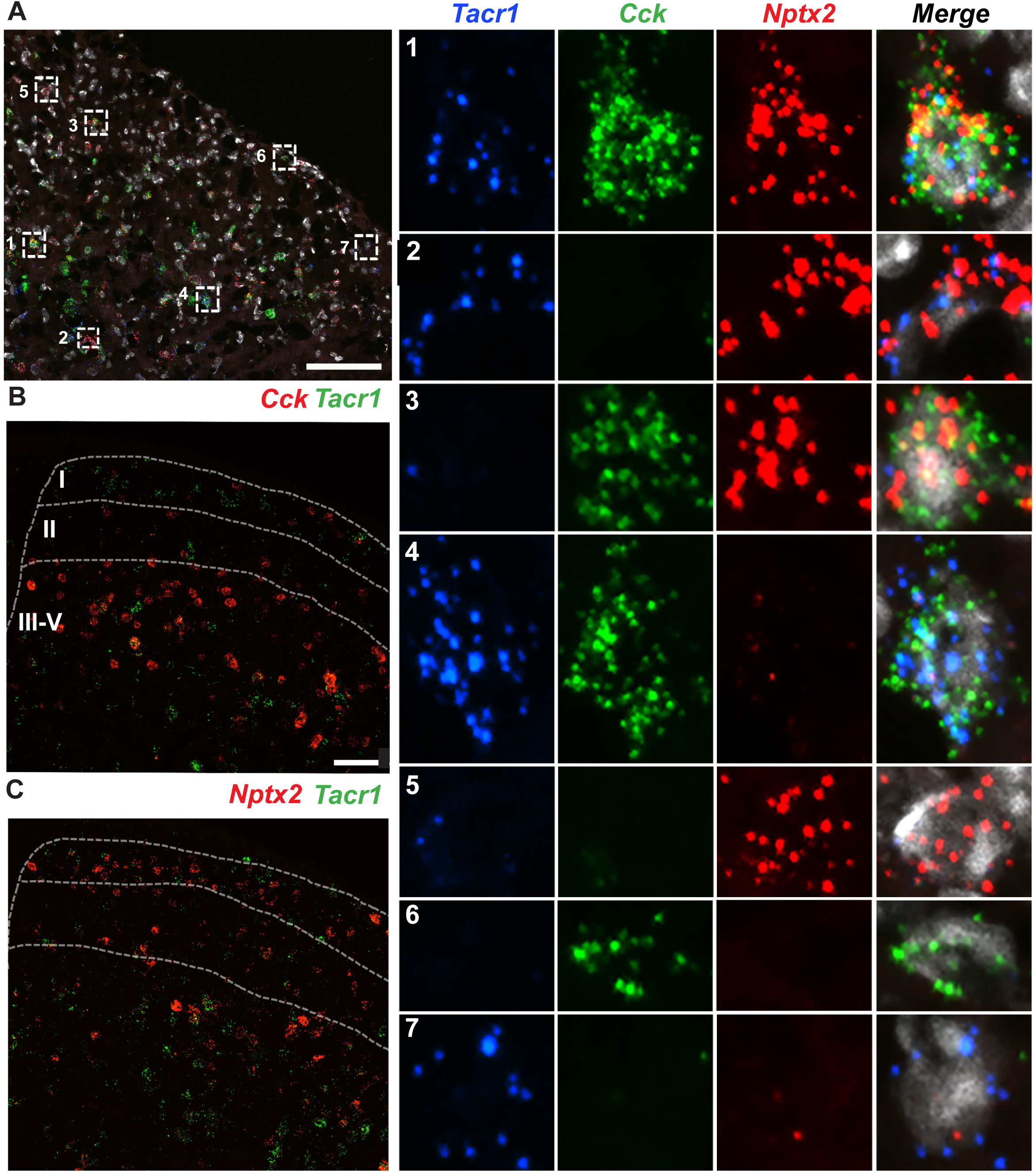
Molecular heterogeneity of dorsal horn *Tacr1*-expressing neurons. **(A)** Representative section of lumbar spinal cord after triple ISH for *Tacr1* (blue), *Cck* (green), *Nptx2* (red), and DAPI (white). Insets show examples of enlarged single neurons with different gene expression combinations, including triple-labeled **(A,1)**, double-labeled **(A, 2-4)** and single-labeled **(A, 5-7)** cells. **(B-C)** Representative sections of lumbar spinal cord after double ISH for *Tacr1* (green) and **(B)** *Cck* (red) or **(C)** *Nptx2* (red). Scale bars, 100μm.

We also investigated whether *Cck and Nptx2* defined non-overlapping subpopulations of NK1R-expressing neurons. Using triple fluorescent ISH for *Tacr1, Cck, and Nptx2* we, in fact, observed neuron subtypes that express every combination of the genes (**Fig. 3A**). Specifically, some neurons triple-label for *Tacr1, Cck, and Nptx2* (**Fig. 3A,1**); others coexpress two of the genes (**Fig. 3A, 2-4**), and others only express one of the three (**Fig. 3A, 5-7**). Based on these results, we conclude that there are at least 4 subsets of *Tacr1*-expressing neurons: *Tacr1*+*Cck*+*Nptx2*+ **(Fig. 3A1)**, *Tacr1*+*Cck*+*Nptx2-* **(Fig. 3A4)**, *Tacr1*+*Cck-Nptx2*+ **(Fig. 3A2)**, *and Tacr1*+*Cck-Nptx2-* cells **(Fig. 3A7)**. Although the NK1R is expressed by some interneurons (Al Ghamdi et al., 2009), it is nevertheless considered a reliable marker of projection neurons. Based on our new results, we conclude that the NK1R-expressing projection neuron population is, in fact, not at all homogeneous. Rather, *Tacr1* is but one marker of a molecularly heterogeneous population of projection neurons.

Recently, several studies reported that substance P (*Tac1*), which targets the NK1R, is also expressed by a subset of the NK1R-expressing projection neurons (Cameron et al., 2015; Huang et al., 2019). We confirmed these findings. Thus, our RNA-seq and qPCR analysis not only identified *Tac1* as an NK1R projection neuron marker (∼4-fold enrichment by RNA-seq), but using ISH, we also demonstrated significant coexpression of *Tac1* and *Tacr1* mRNA in a subset of neurons labeled with Retrobeads (**Supplementary Fig. 3A**). As to other reported genes expressed in projection neurons, Häring et al. (2018) delineated several that cluster with *Tacr1* in the dorsal spinal cord, including *Lypd1* and *Elavl4*. We confirmed these findings by RNA-seq (**Fig. 1H**) and by ISH in spinoparabrachial projection neurons (**Supplementary Fig. 3B**).

### Pain and itch-provoking stimuli engage subsets of molecularly-defined projection neurons

We next used *Fos* mRNA expression to monitor the responsiveness of the retrogradely labeled (Retrobead) projection neuron subsets to an algogenic/painful stimulus (submerging one hindpaw in 50oC water for 30s) or a pruritic/itch-provoking stimulus (cheek injection of chloroquine, CQ: 5.0 µg/µl). To prevent movement and scratching-provoked *Fos*, the mice were anesthetized throughout the experiment. Twenty-minutes after stimulation the mice were killed and we performed triple label ISH in lumbar spinal cord and TNC. The total cell counts (N) for each experiment were obtained by combining data from 2-4 mice with 3-6 sections each, and included Retrobead-labeled cells from superficial and deep laminae of the dorsal horn and TNC, as well as the LSN.

### Pain-relevant projection neurons (***Fig. 4***)

As expected *Fos* induction was most pronounced ipsilateral to the stimulus **(Supplementary Fig. 4**). As expected, we recorded the greatest number of heat-induced *Fos*+ neurons in superficial laminae (I/II) and *Cck*+ projection neurons in laminae III/IV (**Fig 3B**), and in both regions we observed double-labeled *Fos* and *Cck*-positive projection neurons (**Fig. 4A**). On average, about 33% of all projection neurons in lumbar cord were *Cck*-positive, and 9% expressed *Cck* and were noxious-heat responsive, i.e. *Fos positive* (**Fig. 4B**) indicating that ∼27% of the *Cck*-projection neurons responded to noxious heat. For the *Nptx2* population, we observed 41% of projection neurons that were *Nptx2*+ and 21% that coexpressed *Nptx2* and *Fos* after noxious heat stimulation (**Fig. 4C,D**), indicating that ∼50% of the *Nptx2*-expressing projection neurons respond to the noxious heat stimulus. As both *Nmb* (**Fig. 4E, F**) and *Crh* (**Fig. 4G, H**) have highly restricted expression patterns, as expected, we only recorded a few cells per section that were positive for either gene. In this limited number, we found that 36% of the projection neurons were *Nmb*-positive and 10% of all projection neurons were both *Nmb* and *Fos* expressing, indicating that approximately 27% of the limited number of *Nmb*+ projection neurons are “pain” responsive. Finally, for the *Crh* population, we recorded 26% of projection neurons expressing the gene, and 12% of projection neurons that expressed *Crh* responded to noxious heat, suggesting that roughly 50% of *Crh*-positive projection neurons respond to noxious heat.

**Figure 4:**
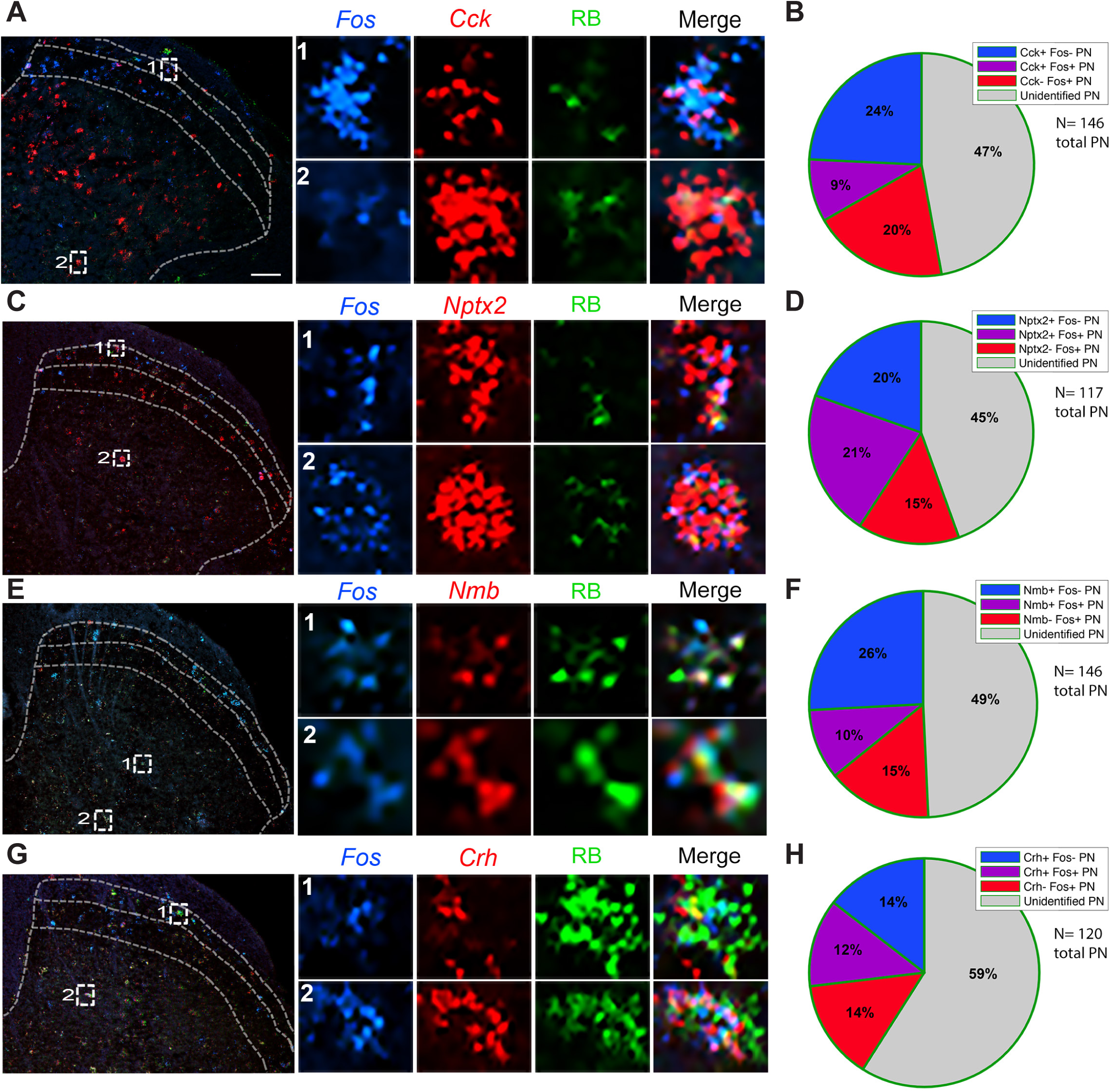
A subset of molecularly-defined projection neurons responds to noxious heat. **(A, C, E, G)** Representative images of lumbar spinal cord illustrate Retrobead-labeled spinoparabrachial neurons (green) co-expressing *Cck* **(A)**, *Nptx2* **(C)**, *Nmb* **(E)**, or *Crh* **(G)** (red), and the *Fos* immediate-early gene (blue), after hindpaw stimulation with noxious heat (50°C). Insets show enlarged examples of triple-labeled cells. **(B, D, F, H)** Pie charts illustrate percentages of projection neurons that express *Fos* after noxious heat stimulation, percentage of projection neurons that express *Cck* **(B)**, *Nptx2* **(D)**, *Nmb* **(F)**, or *Crh* **(H)**, as well as the percentage of projection neurons that express both *Fos* and the marker gene. Quantification includes data from 2-4 mice per gene and 3-6 lumbar spinal cord sections per mouse. Scale bars, 100μm.

### Itch-relevant projection neurons (***Fig. 5***)

We found that 24% of the projection neuron population in the TNC expressed *Cck and* 29% of these *Cck*+ projection neurons were activated by pruritic stimulation; many were in deep dorsal horn. In other words, only 7% of all projection neurons coexpressed *Fos* and *Cck* (**Fig. 5B**). We found that 33% of TNC projection neurons express *Nptx2*; 36% of these projection neurons responded to chloroquine. The latter neurons comprised 12% all projection neurons (**Fig. 5D**). In contrast to the chloroquine responsive *Cck*+ population, the *Nptx2*+ responsive neurons predominated in lamina I. As for the *Nmb* population of TNC projection neurons, we found that 25% of projection neurons express *Nmb* and 8% coexpressed *Nmb* and *Fos*, i.e., 32% of *Nmb*-expressing projection neurons responded to chloroquine (**Fig. 5F**). Lastly, we observed that only 14% of the TNC projection neurons express *Crh* and 4% coexpresed *Crh* and *Fos*, indicating that 28% of *Crh* positive projection neurons in the TNC respond to the pruritogen (**Fig. 5H**). To summarize, we report here that approximately one third of each dorsal horn and TNC population of *Cck-, Nptx2*-, *Crh-*, and *Nmb-*expressing projection neurons responds to heat and/or chloroquine. The extent of convergence is addressed in the next set of experiments.

**Figure 5:**
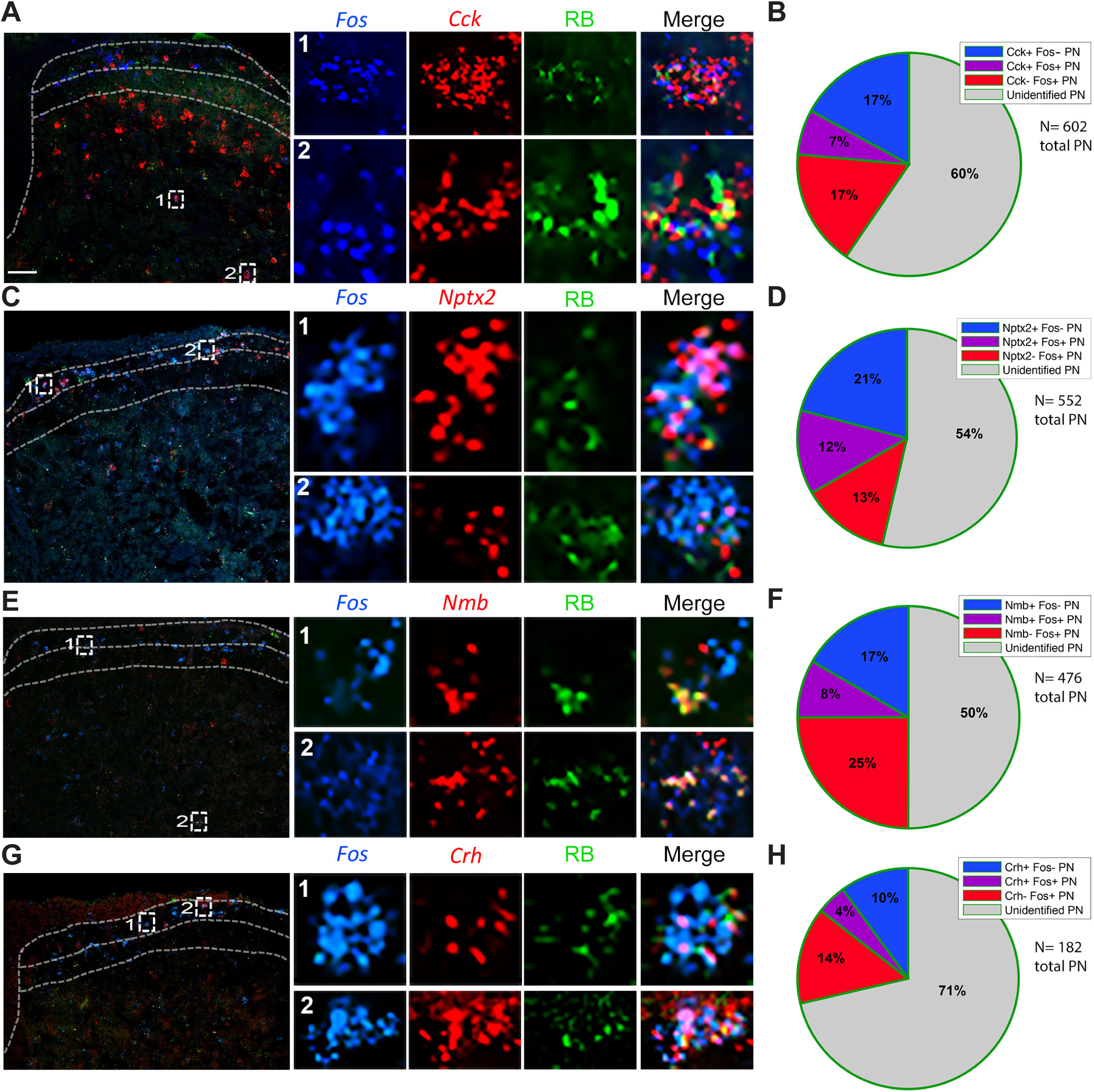
A subset of molecularly-defined projection neurons responds to pruritic (chloroquine) stimulation. **(A, C, E, G)** Representative images of trigeminal nucleus caudalis illustrate Retrobead-labeled trigeminoparabrachial neurons (green) co-expressing *Cck* **(A)**, *Nptx2* **(C)**, *Nmb* **(E)** or *Crh* **(G)** (red) and the *Fos* immediate-early gene (blue), after chloroquine injection into the cheek. Insets show enlarged examples of triple-labeled cells. **(B, D, F, H)** Pie charts illustrate percentages of projection neurons that express *Fos* after chloroquine injection, percentage of projection neurons that express *Cck* **(B)**, *Nptx2* **(D)**, *Nmb* **(F)**, or *Crh* **(H)**, as well as the percentage of projection neurons that express both *Fos* and the marker gene. Quantification includes data from 2-4 mice per gene and 3-6 lumbar spinal cord sections per mouse. Scale bars, 100μm.

### Pain- and itch-provoking stimuli converge upon projection neurons

Here, we used the TRAP2 (Targeted Recombination in Active Populations) mice (Allen et al., 2017) to visualize neurons activated by two stimuli separated in time (**Fig. 6, Supplementary Figs. 5,6**). After exposure to a stimulus delivered within a specific time frame, neurons that are activated by that stimulus) permanently express tdTomato (tdT), a reporter that is expressed in a Cre- and Fos-dependent manner. In these studies, 3h after injecting 4-OH-tamoxifen (100 mg/kg), we anesthetized the mice and then applied a noxious heat stimulus (submerging the mouse hindpaw 5 times in 50°C water for 30s, at 30s intervals). One week later, in the absence of 4-OH-tamoxifen, we injected chloroquine (200 µg) into the same hindpaw. These studies were also performed under continuous anesthesia, to prevent scratching-evoked Fos, These mice were killed 90 minutes later and lumbar spinal cord was processed for immunocytochemical demonstration of the Fos protein. Importantly, these studies were performed in mice that were injected with Fluoro-Gold (FG) into the LPb one week prior to the first stimulation, which allowed for analysis of convergent activation in projection neurons.

**Figure 6:**
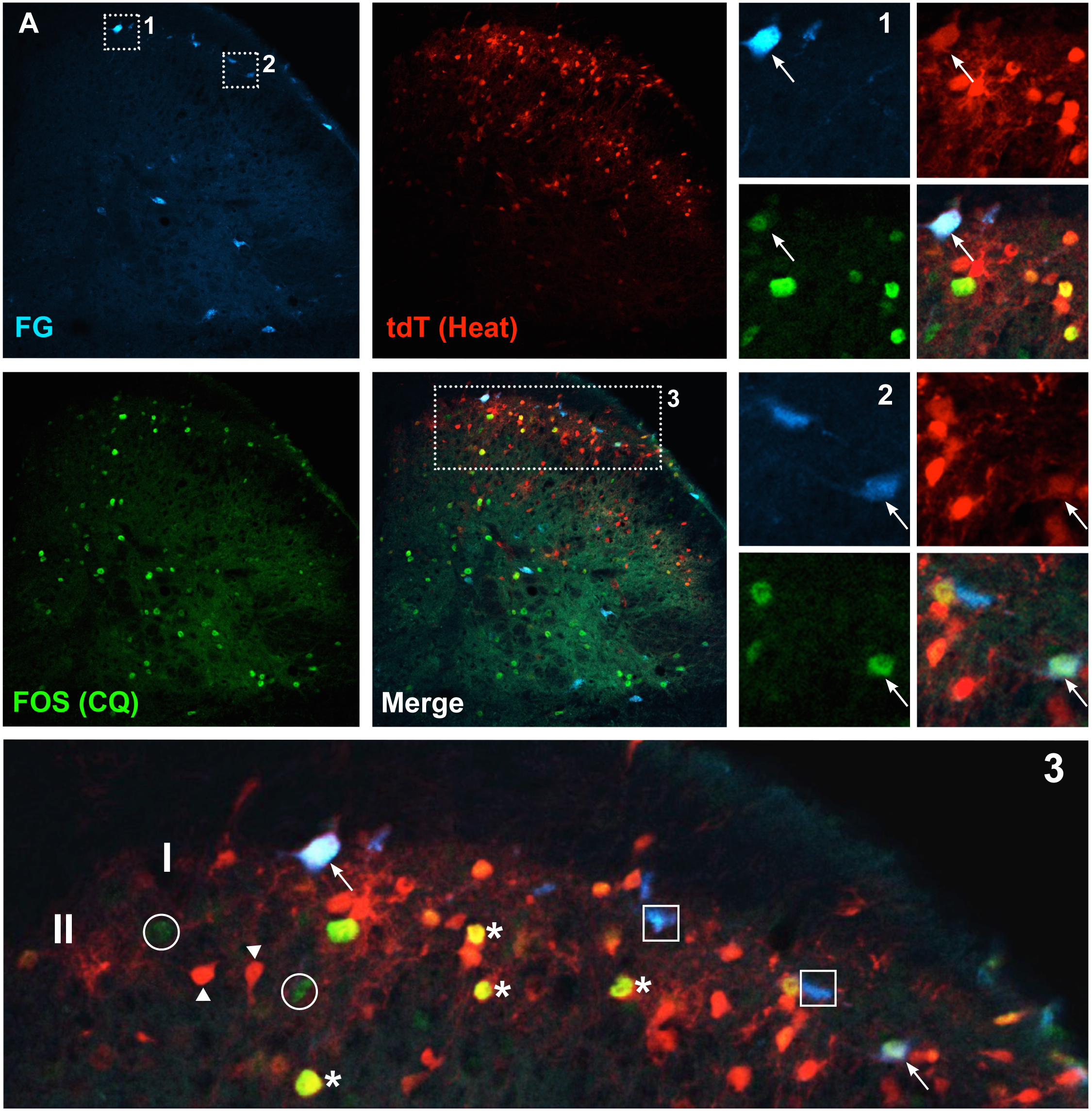
Retrogradely labeled projection neurons respond to both pain (noxious heat) and itch (choroquine)-provoking stimulation. Representative images of lumbar spinal cord from TRAP2-tdTomato mice illustrate Fluorogold-labeled spinoparabrachial projection neurons (FG; blue), heat-activated tdTomato (tdT; red) or chloroquine (CQ)-activated Fos (green) positive neurons. Insets show enlarged examples of single-labeled (circles, squares and arrowhead), double-labeled (stars) and triple-labeled (arrows) neurons. Scale bars, 100μm.

As expected, tdT fluorescence (endogenous) and Fos immunoreactivity were more pronounced in the dorsal horn ipsilateral to the stimulus (**Supplementary Fig. 5**). Demonstrating convergence, we found that many of the projection neurons were both tdT- and Fos-positive (**Fig. 6A-1, −2, −3, Supplementary Fig. 6C**). These triple-labeled cells (i.e., FG+, Fos+, tdT+) predominated in lamina I. By contrast, double-labeled interneurons, i.e., FG-, Fos+, tdT+, were prevalent in lamina II. Other projection neurons in laminae I and III/IV responded only to the noxious (heat) stimulus, i.e., were tdT-positive and Fos-negative (**Supplementary Fig. 6B**), while others responded only to the pruritogen (i.e., were tdT-negative and Fos-positive (**Supplementary Fig. 6A**). Taken together, these studies indicate that subsets of projection neurons transmit both pain and itch relevant messages to the brain, intermingled with others that are more selective, possibly exclusively responsive, to one or the other stimulus.

### Molecularly distinct projection neuron populations are functionally heterogeneous

To integrate the molecular heterogeneity findings demonstrated above with the functional heterogeneity revealed in the TRAP2 studies, we next performed a highly multiplexed, albeit non-quantitative ISH analysis, on lumbar dorsal horn tissue from TRAP2 mice stimulated with both pain- and itch-provoking stimuli (**Fig. 7, 8, Supplementary Fig. 7**). In these experiments, after the 4-OH-tamoxifen, we first “TRAP”ed itch-responsive neurons by injecting chloroquine into the hindpaw and one week later we stimulated pain-responsive neurons by submerging the same hindpaw in 50°C water. To identify projection neurons, these studies were performed in mice that were injected with Retrobeads into the LPb one week prior to the chloroquine. We killed the mice 15-30 minutes after the second stimulus and performed multiplexed ISH for *tdT* (itch-responsive neurons), *Fos* (pain-responsive neurons), *Tacr1, Cck, Nptx2*, and *Crh* (**Fig. 7, 8, Supplementary Fig. 7**). In both the superficial (**Fig. 7**) and deep dorsal horn (**Fig. 8A-G**), as well as the LSN (**Fig. 8H-I**) we observed projection neurons that responded only to the pain-provoking stimulus, i.e., are *Fos*-positive and *tdT*-negative. Some projection neurons responded to both the pain- and itch-provoking stimuli (**Fig 7H-J, Fig. 8A-C,H**) and others did not respond to either (**Fig. 7K-L, Fig. 8F**). Less frequently, we observed neurons that responded only to the itch-provoking stimulus (**Fig. 7G**). Importantly, each of these activated projection neurons expressed varying combinations of the marker-genes (**Fig 7A-F, Fig. 8D-E,I**). As expected most of the projection neurons express the NK1R and one or more of the marker genes, however, by integrating the TRAP2 analysis we now demonstrate that these neurons are also functionally heterogeneous, including itch and pain-responsive subsets. Lastly, we found that the non-NK1R-expressing projection neuron are also molecularly diverse, but in this limited population we did not observe examples of convergence of pain and itch inputs on to these cells (**Fig. 9**). Overall, we conclude that despite limited evidence for functional labeled lines, the molecular diversity of the projection neurons that respond to both pain and itch-provoking inputs provides a basis for the transmission of distinct functional messages to the brain.

**Figure 7:**
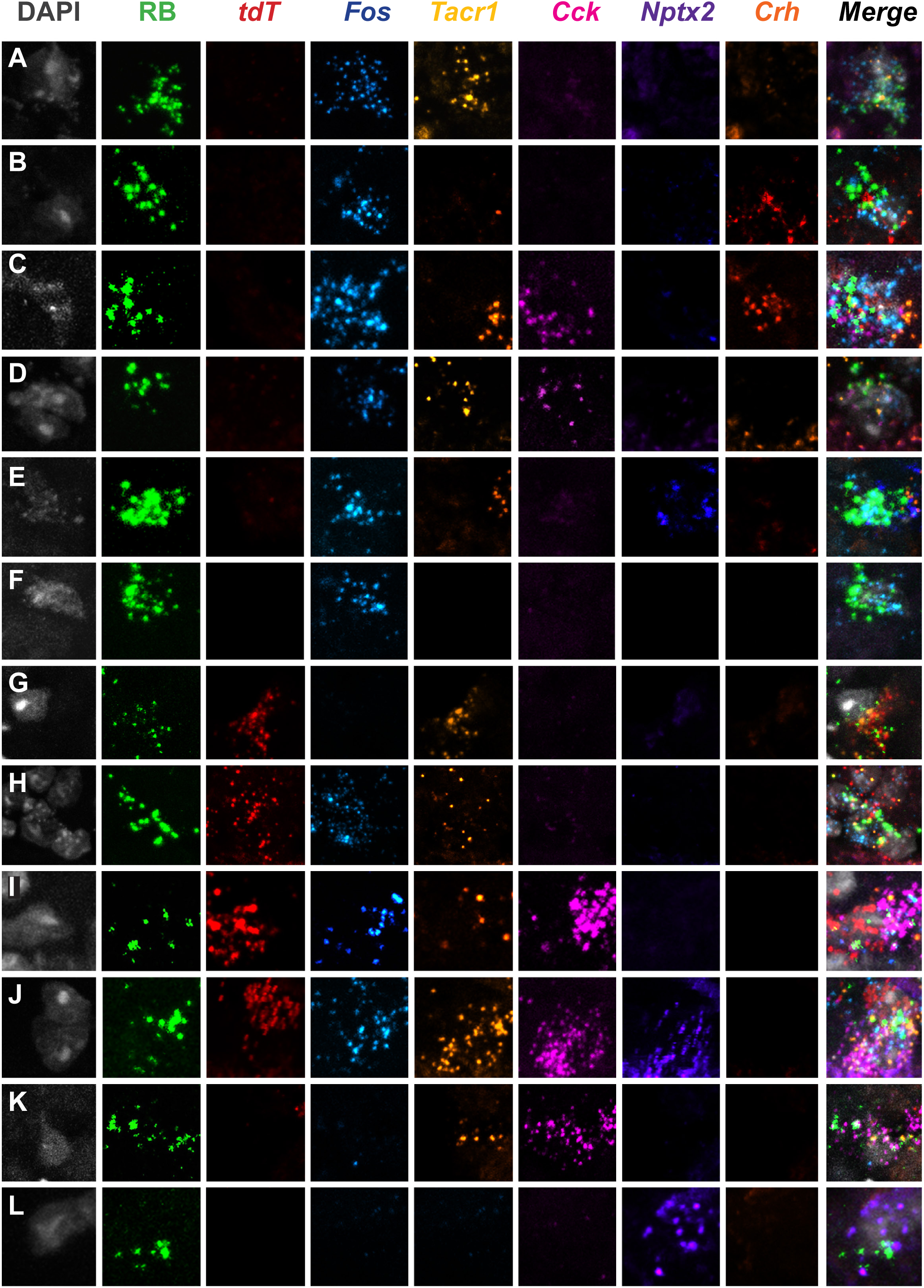
Highly multiplexed *in-situ* hybridization reveals subsets of molecularly and functionally diverse projection neurons in the superficial dorsal horn. **(A-L)** Representative superficial dorsal horn section illustrates Retrobead (RB)-labeled projection neurons from heat (*Fos*) and chloroquine (*tdT*)-stimulated TRAP2 mice. A subset of projection neurons are activated by heat (50°C), but not chloroquine **(A-F)**; other subsets are activated by chloroquine only **(G)**, both heat and chloroquine **(H-J)**, or neither stimulus **(K-L).** Each functionally-defined subset includes projection neurons that express varying combinations of the molecular markers *Tacr1, Cck, Nptx2*, and *Crh*.

**Figure 8:**
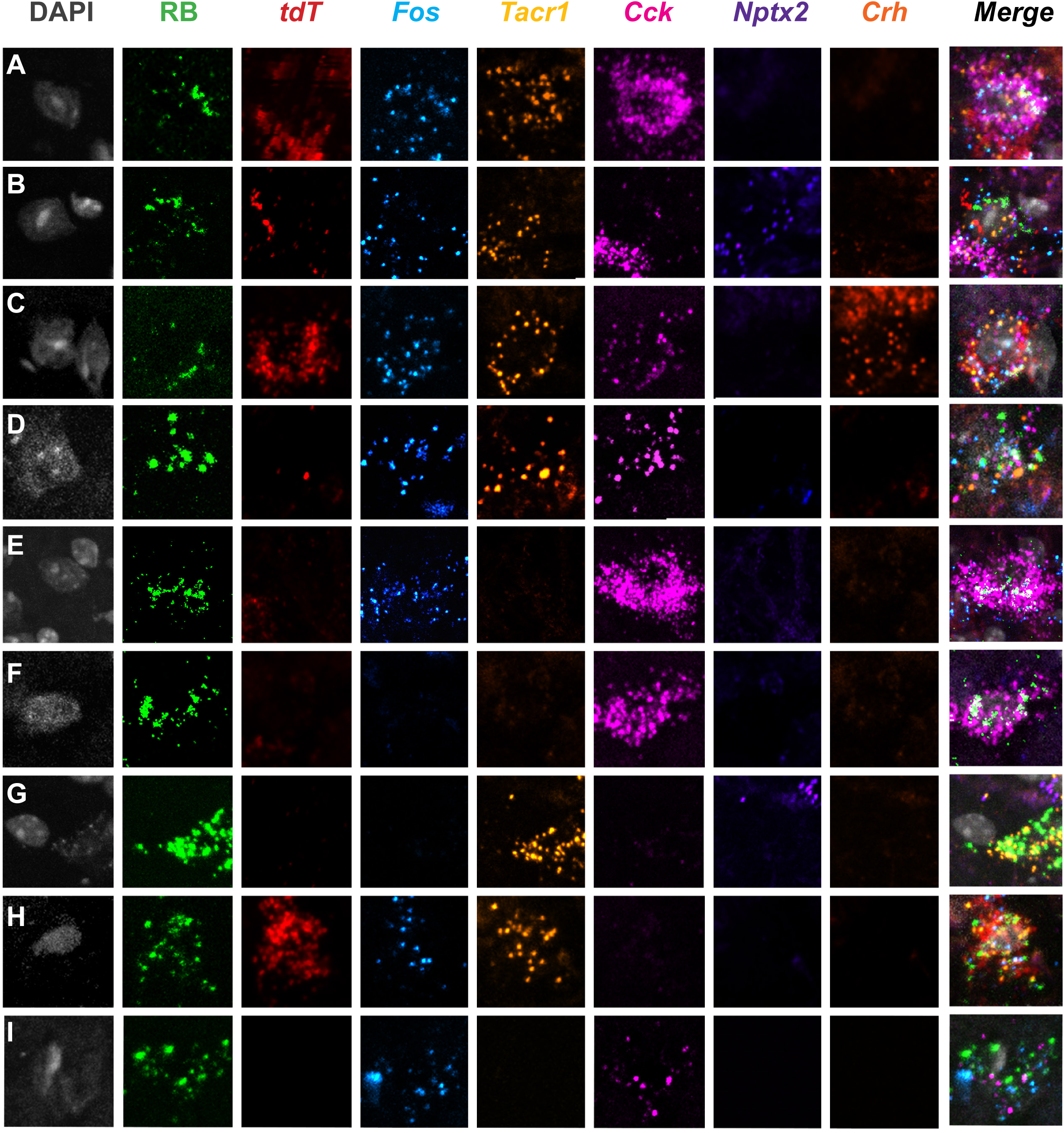
Highly multiplexed *in-situ* hybridization reveals subsets of molecularly and functionally diverse projection neurons in the deep dorsal horn and lateral spinal nucleus. Representative Retrobead (RB)-labeled projection neurons in deep dorsal horn **(A-G)** and lateral spinal nucleus **(H-I)** of TRAP2-stimulated mice. A subset of projection neurons **(A-C, H)** are activated by both 50°C heat (i.e., *Fos*+) and chloroquine (i.e., *tdT*+). Other subsets are activated by heat only **(D-E, I)** or by neither stimulus **(F-G).** Each subset includes projection neurons that have varying combinations of the molecular markers: *Tacr1, Cck, Nptx2*, and *Crh*.

**Figure 9:**
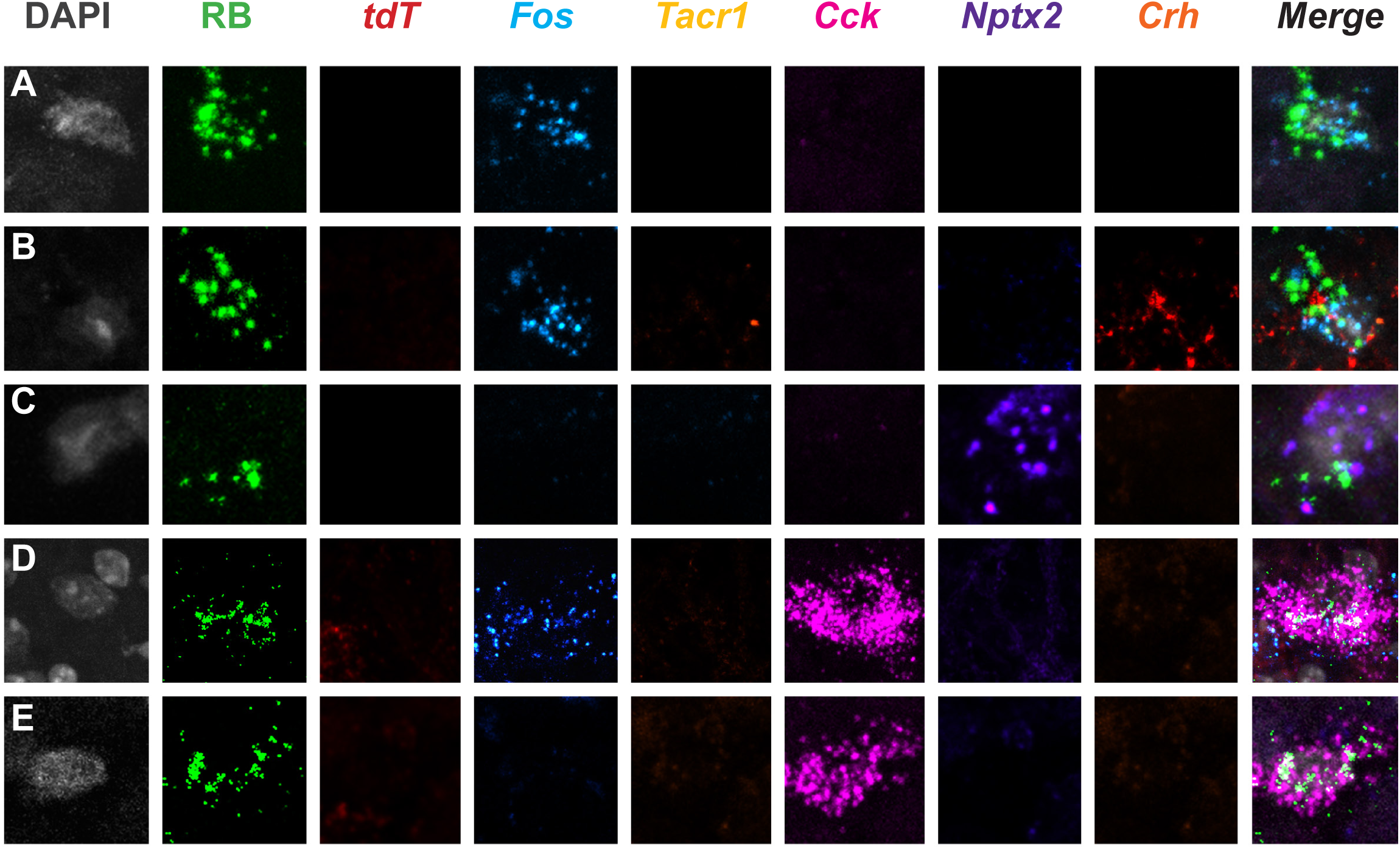
Projection neurons that do not express the NK1R are also molecularly heterogeneous. Representative Retrobead (RB)-labeled projection neurons in superficial **(A-C)** and deep dorsal horn **(D-E)** of TRAP2 mice that are negative for *Tacr1* express different combinations of *Cck, Nptx2*, and *Crh*. The illustrated projection neurons respond either only to heat (50oC; i.e., *Fos* mRNA+; **A, B, D**) or neither to heat nor chloroquine (*Fos* mRNA- and *tdT* mRNA-negative; **C, E**).

## DISCUSSION

There is general agreement that primary sensory neurons and dorsal horn interneurons are heterogeneous, responding, to different degrees, selectively to pain- and or itch-provoking stimuli. Of course, the brain generates functionally distinct percepts based on the information transmitted by the projection neurons, not by primary sensory neurons or dorsal horn interneurons. Surprisingly, therefore, despite considerable evidence that the projection neurons that transmit pain and itch are diverse in regards to location, projection targets, morphology, and electrophysiological properties, reference is often made to a rather homogeneous molecular and functional population of projection neurons. Based on the present ribosomal profiling of the LPb-projecting neurons we conclude that there are, in fact, molecularly and functionally distinct subpopulations of projection neurons that can be distinguished based on gene expression, spatial location in the dorsal horn and TNC, and responsiveness to pain and/or itch-provoking stimulation. To our knowledge, the current study provides the first comprehensive investigation into the molecular profiles and functional properties of projection neuron subtypes.

### Technical considerations

As noted above, we designed the initial filtering steps to identify neuronal genes that are expressed by projection neurons. Given the nature of the input samples, which included all spinal cord and TNC cells, glial and neuronal, we recognize that many of the RNA-seq hits, although clearly enriched in neurons relative to glia, also included genes expressed by interneurons. However, the subsequent filter steps, taken together with a comparison of results with the Häring et al. (2018) database, directed our attention to genes that, although not exclusively expressed in projection neurons, unequivocally defined subtypes. Furthermore, because there is considerable variability in retrograde labeling, it was never our intention to quantify the percentage of projection neurons that express each marker. Rather these filtering steps identified genes that informed the subsequent *in situ* and functional studies that demonstrated molecular and functional heterogeneity of the projection neurons.

The discrepancy between mRNA signal, as visualized by ISH, and protein level detected by IHC, raises interesting questions. That is, consistently, we detected significantly greater numbers of neurons expressing a given message rather than its translated protein. This is particularly true for the NK1R, where we recorded large numbers of *Tacr1* mRNA positive neurons in laminae III/IV, but few immunoreactive cells. This difference between mRNA and detectable protein could either be due to increased sensitivity of the ISH compared to immunohistochemistry or it could reflect a true difference between the rates of gene transcription and translation. Which technique better illustrates the functional populations of NK1R-expressing neurons remains to be determined. It follows that the extent to which we conclude that there is coexpression of markers indicative of molecular heterogeneity and convergence of inputs to individual projection neurons may be underestimated (using immunohistochemistry) or overestimated (using ISH). In addition, the retrograde viruses and tracers that we used may not have captured the entire projection neuron population. Lastly, conclusions based on the TRAP2 studies are highly dependent on the dose of 4OH-tamoxifen and timing of the two different stimuli.

### NK1R is not the ideal projection neuron marker

A common trend in the literature is to equate “pain” and “itch” projection neurons with NK1R expression. Although Todd and colleagues determined that 80% and ∼90% of lamina I projection neurons in the rat (Marshall et al., 1996; Spike et al., 2003) and mouse (Cameron et al., 2015), respectively, express the NK1R, less than half of deep dorsal horn projection neurons express the receptor (Cameron et al., 2015; Marshall et al., 1996). Studies that used the NK1R to define the functional contribution of those projection neurons also have limitations. For example, Mantyh et al. (1997) and Carstens et al. (2010) found that ablating NK1R-expressing neurons reduces behaviors indicative of both pain and itch. At first glance this suggests that pain and itch inputs converge on NK1R-expressing projection neurons. Based on our present findings, however, it is clear that there are several subpopulations of NK1R-expressing neurons that transmit modality specific information to the brain. We conclude, therefore, that the NK1R is not the ideal marker to interrogate specificity at the level of the projection neurons.

### Non-NK1R projection neurons

Although most superficial dorsal horn projection neurons express the NK1R, about 10-20% of the lamina I projection neurons do not (Todd, 2010). The same is true for the majority of projection neurons in laminae III-V, which despite considerable literature implicating them in pain processing, have largely been ignored (Wercberger and Basbaum, 2019). In the present study, we compared RNA-seq data of projection neurons from wildtype mice (i.e., all LPb-projecting neurons of the spinal cord dorsal horn and TNC) with those from NK1R-Cre mice (i.e., NK1R-expressing projection neurons). Using differential analysis, we generated a list of genes that are significantly enriched in the PN dataset and depleted in the NK dataset. We hypothesize that these genes are selectively enriched in the non-NK1R-expressing projection neurons. Although pursuit of these genes is not the subject of the present study, we present them as a resource for future investigation. Additionally, our RNA-seq data provide a preliminary confirmation of a recent identification of Gpr83 expression in the non-NK1R-expressing projection neurons (David Ginty, personal communication). Specifically, although modest, we found a ∼1.7-fold enrichment of Gpr83 in the PN dataset, but no enrichment in the NK dataset (**Supplementary Files 1,2**). On the other hand, we did not find enrichment of two other reported markers of the non-NK1R-expressing projection neurons, namely the glycine α1 (GlyRα1) and GluR4 AMPA receptor subunits (Polgár et al., 2008; Puskár et al., 2001). Finally, our studies using Hiplex ISH revealed examples of molecularly diverse non-NK1R-expressing projection neurons. In contrast to our findings of algogen and pruritogen convergence on NK1R projection neurons, however, we did not find comparable convergence in what admittedly was a limited sample of non-NK1R projection neurons. For the same reason we are hesitant to conclude that there is greater specificity of modality transmission along the non-NK1R population.

### Molecular properties of projection neurons vs existing dorsal horn neuronal transcriptomics

Recent single-cell and single-nucleus RNA sequencing studies reported on the molecular heterogeneity of spinal cord neurons. Sathyamurthy et al. (2018) identified 16 dorsal horn excitatory neuron types, but interestingly, they concluded that none are defined uniquely by NK1R expression. In fact, they found expression of *Tacr1* across several clusters and made no mention of projection neurons. In contrast, Häring et al. (2018) delineated 15 dorsal excitatory neuron categories and, by integrating a retrograde tracing approach, concluded that spinoparabrachial neurons are concentrated within one of the 15 clusters of excitatory neurons (Glut 15). Our study differs considerably from these studies in approach, experimental procedures, and most importantly, in conclusions. Thus, to specifically characterize the molecular heterogeneity of projection neurons, we used bulk profiling of isolated projection neuron ribosomes from dorsal spinal cord and TNC tissue. Not surprisingly, we concur with Häring et al.’s identification of Glut 15, one of the clusters defined by NK1R enrichment, as a projection neuron population. Specifically, as predicted by Häring et al. we identified *Crh, Lypd1* and *Elavl4* in projection neurons, all of which are included in the Glut 15 cluster. Importantly, although Häring et al. proposed *Lypd1* as a novel marker for projection neurons, because *Lypd1* is expressed in almost 95% of NK1R projection neurons, it cannot be used to define subtypes. In contrast, we identified more discrete markers of projection neurons and interestingly, these markers belong to excitatory populations other than Glut15. For example, our finding of *Cck*+, *Nptx2*+ and *Nmb*+ projection neurons demonstrates that there are spinoparabrachial neurons within the Glut 2, 3, 7, 9, and 11-14 excitatory clusters. On the other hand, as many of these marker genes are also expressed in sensory neurons and dorsal horn interneurons, we emphasize that they cannot be used as sole markers of projection neurons. Future studies of their relative functional will require complex intersectional approaches.

### Specificity versus convergence and the generation of pain and itch percepts

Using techniques that selectively stimulate or ablate subsets of neurons (e.g. DREADDS, intersectional knockouts, optogenetics) recent studies have provided considerable evidence for a labeled-line view of pain and itch transmission, at least at the level of primary sensory neurons and dorsal horn interneurons (Koch et al., 2018; Kupari et al., 2019; LaMotte et al., 2014; Peirs and Seal, 2016). On the other hand, with some exceptions (Craig et al., 2001; Dostrovsky and Craig, 1996), electrophysiological data overwhelmingly suggest polymodality at all levels of the neuraxis (Sun et al., 2017). The latter implies that pain- and itch-provoking inputs either converge on modality-indiscriminate circuits, a conclusion that ignores results from many studies, or that a population code is in effect, one that results from cross talk among different labeled-lines. In fact, with regard to pain, itch, and thermal processing, several groups described combinatorial coding at the level of the DRG (Wang et al., 2018) and spinal cord (Ran et al., 2016; Sun et al., 2017). Indeed, Sun et al. (2017) reported that GRP interneurons, generally considered to exclusively contribute to the processing of itch messages, also contribute to pain processing depending on noxious stimulus intensity. Our findings are consistent with a population code model of pain and itch processing. We speculate that a similar code underlies the processing of pain and itch messages by the molecularly diverse population of projection neurons that receive convergent modality inputs. To what extent a labeled line component integrates with the output of projection neurons that use a population code in the generation of pain and itch and most importantly how the brain interprets these messages remains to be determined.

## METHODS

### Animals

Animal experiments were approved by the UCSF Institutional Animal Care and Use Committee and were conducted in accordance with the NIH Guide for the Care and Use of Laboratory animals. Male and female wild type C57BL/6J mice were purchased from Jackson Laboratory (Stock #000664). Transgenic mice that express Cre-recombinase in NK1R-expressing neurons were kindly provided by Dr. Xinzhong Dong (Baltimore, USA, see also Green et al., 2019). TRAP2 transgenic mice were purchased from Jackson Laboratory (Stock #030323; see also Allen et al., 2017). All mice used were between 6-12 weeks of age and were housed on a 12 hr light-dark schedule.

### Stereotaxic injections

To tag spinal cord dorsal horn and trigeminal nucleus caudalis (TNC) projection neuron ribosomes for immunoprecipitation, we first anesthetized adult mice with ketamine (60mg/kg) / xylazine (8.0mg/kg). Bilateral craniotomies were made over the lateral parabrachial nucleus (LPb, coordinates: ±1.3 mm from midline, −5.34 mm from Bregma, −3.6 mm from skull), into which we injected 0.5μl of a herpes simplex-based viral vector expressing the large ribosomal subunit protein L10a (gift from Dr. Zachary Knight, UCSF), which is transported retrogradely to the spinal cord and TNC. In the wild type mice, we used a viral vector that expresses a GFP-tagged L10a (HSV-hEF1a-GFP-L10a) and in the NK1R-Cre mice we used a viral vector that expresses Cre-recombinase-dependent HA-tagged L10a (HSV-hEF1a-LS1L-Flag-HA-L10a). After each viral injection, the needle was left in place for 5 minutes before slowly retracting. The skin was sutured and the mice were returned to their cages, administered buprenorphine-SR (0.15 mg/kg, i.p.) and monitored for recovery. Mice were killed 2-3 weeks after surgery and their tissue dissected for immunoprecipitation.

To study the molecular properties of projection neurons, we retrogradely labeled projection neurons of the spinal cord dorsal horn and TNC in wild type mice by making either unilateral or bilateral LPb injections of 0.5 μl of green Retrobeads (Lumafluor), 2% Fluoro-Gold (Fluorochrome), or the HSV-hEF1a-GFP-L10a.

### Immunoprecipitation, RNA and cDNA preparation

To immunoprecipitate tagged ribosomes and their associated mRNA, we followed the protocol described by Ekstrand et al. (2014) with minor modifications. Protein A Dynabeads (100 μl per IP, Invitrogen) were washed twice on a magnetic rack with Buffer A (10 mM HEPES [pH 7.4], 150 mM KCl, 5.0 mM MgCl2, 1.0% NP40), resuspended in Buffer A with 0.1% BSA and then loaded with either anti-GFP polyclonal antibody (25 µg per IP, abcam) or anti-HA-tag monoclonal antibody (2.0 µg per IP, Cell Signaling). Antibody-bead conjugates were mixed at 4°C overnight.

Mice were killed with an overdose of Avertin (2.5%) and dorsal spinal cord and TNC were rapidly dissected in ice-cold Buffer B (1xHBSS, 4.0 mM NaHCO3, 2.5 mM HEPES [pH 7.4], 35 mM Glucose) with 100 μg/ml cycloheximide (Sigma). The dissected pieces were pooled in 2 groups of 4 mice each (1 group of injected experimental mice, 1 group of non-injected controls), transferred to a glass homogenizer (Kimble Kontes 20) and homogenized in 1.5 ml ice-cold Buffer C (10 mM HEPES [pH 7.4], 150 mM KCl, 5.0 mM MgCl2) with 0.5 mM DTT (Sigma), 20 U/μl Superase-In (Invitrogen), 100 μg/ml cycloheximide, and protease inhibitor cocktail (Roche).

Tissue samples were homogenized 3 times at 300 rpm and 10 times at 800 rpm on a variable-speed homogenizer (Glas-Col) at 4°C. Homogenates were transferred to microcentrifuge tubes and clarified at 2,000xg for 10 min at 4°C. 10% IGEPAL CA-630 (NP-40; Sigma) and 1,2-diheptanoyl-sn-glycero-3-phospho-choline (DHPC at 100 mg/0.69 ml; Avanti Polar Lipids) were then added to the supernatant for a final concentration of 30 mM and 1%, respectively. The solutions were mixed and centrifuged again at 20,000xg for 15 min at 4°C. The resulting supernatants were transferred to new tubes and 50 μl of each cleared lysate was mixed with 50 μl Lysis Buffer (0.7 μl β-mercaptoethanol/100 μl Lysis Buffer; Agilent Absolutely RNA Nanoprep Kit) and stored at −80°C for later preparation as Input RNA. The remaining lysates (approximately 1.5 ml) were used for immunoprecipitation.

The beads incubating with GFP or HA antibodies were washed twice in Buffer A before the tissue lysates were added. The GFP and HA IPs ran at 4°C for 5 or 10 min, respectively. Beads were washed 4 times with Buffer D (10 mM HEPES [pH 7.4], 350 mM KCl, 5 mM MgCl2, 1% NP40) with 0.5 mM DTT, 20 U/μl Superase-In Plus and 100 μg/ml cycloheximide. Before removing the last wash solution the beads were transferred to a new tube. After the final wash, RNA was eluted by adding 100 μl Lysis Buffer and purified using the Absolutely RNA Nanoprep Kit (Agilent). cDNA was prepared with the Ovation RNA-seq V2 kit (Nugen) and a portion was set aside for analysis with Taqman qPCR (see below for methods). Libraries for RNA-seq were prepared with the remaining cDNA using the Ovation Ultralow Library System (NuGen).

### RNA yield and quality

RNA yield (ng/μl) and quality (RIN value) were quantified using an Agilent 2100 Bioanalyzer and the Agilent RNA 6000 Pico Kit (Cat No. 5067-1513). IP RNA yields for the GFP IP ranged from 0.1-0.9 ng/μl and yields from the HA IP ranged from 2.5-3.5 ng/μl. Input RNA yield ranged from 50-90 ng/μl. We only analyzed samples with RIN values of 8.4 or greater.

For the Input control, we used total mRNA from homogenized dorsal spinal cord and TNC. Consistently, the RNA yields from the IPs performed on tissue from injected animals were at least one order of magnitude greater than IPs performed on tissue from uninjected controls (data not shown). We only detected 18S and 28S rRNA peaks from the former samples, which indicates that the ribosomal tag was critical to the RNA pull down and that the IPs were specific for projection neurons.

### Quantitative PCR

Dorsal spinal cord and TNC tissue were homogenized and RNA and cDNA were prepared as described above. We quantified mRNA levels with the Bio-Rad CFX Connect System using TaqMan Gene Expression Assay (Applied Biosystems). Taqman data analysis and statistics were performed using Microsoft Excel. All Taqman values were normalized to ribosomal protein Rpl27. Fold enrichment plots from Taqman data were obtained by dividing the IP RNA value for each gene by the Input RNA value (IP/Input). A paired Student’s t test was performed to compare IP and Input RNA.

### Sequencing and bioinformatic analysis

RNA sequencing was performed on an Illumina HiSeq 4000 sequencer using 50 bp single-end reads. For the GFP-IP experiment, 8 samples were sequenced: 4 Immunoprecipitate replicates paired with 4 Input replicates, which were obtained from pooling 4 mice per replicate. For the HA-IP experiment, 6 samples were sequenced: 3 Immunoprecipitate replicates paired with 3 Input replicates, which were also obtained from pooling 4 mice per replicate. RNA-seq data were processed in Galaxy and further analyzed with Microsoft Excel and MATLAB (R2015b). RNA STAR (v 2.6.0b) was used to align the reads. Htseq-count (v 0.9.1) and DESeq2 (v 1.18.1) were used for transcript abundance estimation and differential expression testing, respectively. The UCSC GRCm38 (mm10 build) was used for gene annotation.

### In situ hybridization (ISH): RNAscope multiplex hybridization

To detect and confirm expression of marker genes in spinal cord and TNC tissue we performed fluorescent *in situ* hybridization (ISH) using the RNAscope Multiplex Fluorescent Assay (Advanced Cell Diagnostics, cat no. 320850) and target probes. For the complete list of probes used, see Supplementary File 4. For these experiments, we perfused adult C57BL/6J mice with room temperature 1X phosphate-buffered saline (PBS). The TNC and spinal cord were rapidly dissected out, frozen on dry ice, and kept at −80°. From these tissues, we cut 12 µm cryostat sections and stored the sections at −80°C until use. The day of ISH, frozen sections were immediately fixed at 4°C in 4% formaldehyde for 15 minutes, washed twice in PBS and dehydrated through successive ethanol steps (50%, 70% and 100% ethanol) for 5 minutes each and dried at room temperature (RT). After a 30 minute incubation step with protease IV, sections were washed twice in PBS and incubated at 40°C with RNAscope-labeled probes for 2h in a humidified chamber. Sections were then washed twice in washing buffer and incubated with 4 successive “signal amplifying” solutions at 40°C, for 15 to 30 minutes each. After two washes, the sections were dried and covered with mounting media containing DAPI.

For *Fos* induction studies: Mice were injected two weeks prior to stimulation with HSV-hEF1a-GFP-L10a or green Retrobeads (Lumafluor) into the LPb, to label LPb-projecting neurons. For pruritic stimulation, we injected chloroquine (CQ; 500 μg in 100 μL) into the left cheek, and performed ISH in the TNC for *Fos*, and each marker gene. For noxious heat stimulation, we submerged the left hindpaw in 50°C water for 30s and performed ISH on lumbar spinal cord. In mice in which projection neurons were labeled with HSV-hEF1a-GFP-L10a we also used a *Gfp* probe. All mice were injected i.p. with an anesthetizing dose of Avertin (1.25%) 20-30 minutes before stimulation, and killed 15-30 min after stimulation with a lethal Avertin dose.

### In situ hybridization (ISH): RNAscope Hiplex12 hybridization

To simultaneously detect up to 7 marker genes in spinal cord tissue, we performed fluorescent ISH using the RNAscope Hiplex12 ancillary kit (Cat. No. 324140) assay, which is similar to the protocol used for RNAscope multiplex described above, with the following differences: 1) frozen sections were fixed for 60 rather than 15 minutes at RT; 2) sections were incubated for 2h with all 7 probes at once; 3) all 3 amplifying solutions were incubated for 30 minutes each; 4) after 2 washes, the sections were incubated for 15 min at 40°C with the Hiplex Fluoro T1-T4 solution. After 2 final washes, the sections were incubated with the RNAscope DAPI solution for 30 sec at RT and covered with Prolong Gold Antifade mounting media (Fisher Scientific #P36930). Because only 3 targets can be concurrently visualized, due to a limited number of detection channels available for microscopy (488, 550 and 647 channels), we performed 3 successive rounds of detection/imaging/cleaving on 3 consecutive days, as follows. After the first imaging round, the sections were incubated in 4X SSC until the coverglass easily detached from the slide (up to 24h), briefly washed with 4X SSC and then incubated twice for 15 min at RT with 10% cleaving solution (with 2 washes in between). After 2 rinses in washing buffer, the sections were incubated for 15 min at 40°C with the Hiplex Fluoro T5-T8 solution, washed twice, immediately covered with Prolong Gold Antifade mounting media and imaged. A final round of detection/imaging was performed on the 3^rd^ day as described above, using the Hiplex Fluoro T9-T12 solution. Images from each round were merged together using the RNAscope Hiplex registration software 300065-USM and Photoshop to adjust intensity and contrast.

### TRAP2 assay

TRAP2 mice (Allen et al., 2017) were crossed with mice that ubiquitously express tdTomato (tdT) after tamoxifen-induced Cre recombination (Ai14 mice; Jackson Laboratories, stock #007914), which generates double transgenic TRAP2-tdT mice. Adult TRAP2-tdT mice first received a unilateral injection of the retrograde tracer, Fluoro-Gold (2%) into the LPb, as described above. One week later, the TRAP2-tdT mice received an i.p. injection of 4-hyroxy-tamoxifen (100 mg/kg, dissolved in oil; Sigma). Three hours later, the TRAP2-tdT mice were stimulated, under anesthesia, as follows: one group of mice received a subcutaneous injection of chloroquine (200µg in 25µl) into the hindpaw (CQ group) and in another group the entire hindpaw was dipped 5 times for 30 seconds each, into a 50°C waterbath, with 30 sec between each stimulus (heat group). The stimuli were administered contralateral to the retrograde tracer injections. One week later, both groups of TRAP2-tdT mice were stimulated a second time, again under anesthesia, as follows: using the same protocol, the CQ group received the heat stimulus and the heat group received the CQ stimulus. Ninety minutes later, all mice received an overdose of Avertin (2.5% and then lumbar spinal cord tissue was processed for immunohistochemistry or ISH as described above.

### Imaging and image analysis

All images were taken with an LSM 700 confocal microscope (Zeiss) equipped with 405-nm (5mW fiber output), 488-nm (10mW fiber output), 55-nm (10-mW fiber output), and 639-nm (5-mW fiber output) diode lasers using a 20x Plan Apochromat (20x/0.8) objective (Zeiss). To acquire images, we used Zeiss Zen software (2010). The same imagine parameters were used for all images within an experiment.

For quantitative analysis of transcript expression, we analyzed 12μm sections from 2-4 animals per condition (3-6 sections per animal). Because we performed the pruritic stimulation in the cheek, which allowed for a larger chloroquine volume, in these experiments we analyzed TNC tissue. For the noxious heat stimulation experiments, we analyzed lumbar spinal cord (L3-5). We used a custom MATLAB script to count cells positive for the retrograde-label (Retrobeads or GFP) and subsequently determined the percentage of these cells that were positive for the marker genes and/or *Fos*. For quantitative analysis of the spatial distribution of enriched genes, we used MATLAB to manually draw a border between superficial and deep dorsal horn or TNC and used a custom MATLAB script to count the percentage of projection neurons in each region that express each gene. To calculate the final percentages of gene expression and spatial distribution, we averaged counts and percentages across sections in each animal, and again across animals per experimental group.

## Acknowledgements

This work was supported by: NIH NS14627 (AIB), NSR35097306 (AIB) and NSF Graduate Research Program (RW). We thank Dr. Zachary Knight for providing significant advice in the design of experiments.

## Author Contributions

Conception and design (RW, JMB, AIB); Acquisition (RW, JMB); Analysis and interpretation of data (RW, JMB, JAW, AIB); Writing the manuscript (RW, JMB, AIB).

## SUPPLEMENTARY FIGURES

**Supplementary Figure 1: Validation of RNA purification, sequencing and Volcano plots of enriched and depleted genes in LPb-projecting neurons**

**(A)** Bioanalyzer traces of projection neuron IPs from virus injected (top) and control (bottom) animals, for PN (HSV-GFPL10 injected, left) and NK (HSV-HAL10 injected, right) experiments. RIN, RNA Integrity Number. FU, Fluorescence intensity. **(B)** qPCR analysis showing average fold enrichment of *Gfp* mRNA in IP relative to Input from PN experiments. **(C)** qPCR analysis showing average fold enrichment of *Tacr1* mRNA in IP relative to Input from NK experiments. **(D)** qPCR analysis of average glial cell depletion in IPs relative to Input samples for PN and NK experiments. **(E)** qPCR analysis of enriched genes from PN RNAseq dataset **(F)** qPCR analysis of enriched genes from NK dataset **(B-F)** Data are normalized to Rpl27. n = 3 libraries. *p < 0.05 **p < 0.005 ***p < 0.0001 (**G,H**) RNAseq analysis represented as volcano plots for the PN **(G)** and NK experiments **(H)**. Insets show enlarged area with genes of interest highlighted in yellow. Genes significantly changed in both PN and NK datasets are indicated in red, genes significantly changed PN but not NK are shown in green, and genes significantly changed in NK but not PN are shown blue. P < 0.05.

**Supplementary Figure 2: Novel genes that mark subpopulations of NK1R-expressing cells**

Representative sections from lumbar spinal cord showing coexpression of *Cck* **(A)**, *Nptx2* **(B)**, *Nmb* **(C)**, or *Crh* **(D)** mRNA (red) with *Tacr1* mRNA (green) in both superficial and deep dorsal horn. Insets show magnified examples of single-labeled cells for each individual marker gene, or for *Tacr1*, as well as double-labeled cells, in superficial (middle panels) and deep (right panels) dorsal horn. Scale bars, 100μm.

**Supplementary Figure 3: *Tac1, Lypd1* and *Elavl4* are expressed by subsets of NK1R-expressing neurons**

**(A)** Representative section illustrates co-expression of *Tac1* (red) and *Tacr1* (blue) mRNA in Retrobead-labeled trigeminoparabrachial neurons (green). **(B)** Representative section illustrates co-expression of *Lypd1* (blue), *Elavl4* (green) and *Tacr1* (red) mRNA in lumbar dorsal horn neurons. Insets in **A** and **B** show enlarged examples of triple-labeled cells. Scale bars, 100μm.

**Supplementary Figure 4: Algogen (heat) and pruritogen (chloroquine) stimulation evokes asymmetrical *Fos* mRNA in superficial dorsal horn and TNC**

**(A)** Representative sections of lumbar spinal cord illustrate ipsilateral *Fos* expression in sDH in response to noxious heat (50°C). **(B)** Representative sections of TNC illustrate ipsilateral *Fos* expression in response to chloroquine. sDH: superficial dorsal horn; dDH: deep dorsal horn; LSN: lateral spinal nucleus; SC: spinal cord; TNC: trigeminal nucleus caudalia. Scale bars, 100μm.

**Supplementary Figure 5: Permanent labeling of transiently activated spinal cord neurons via the TRAP2 (targeted recombination in active populations) assay** Representative sections of lumbar spinal cord from TRAP2-tdTomato mice illustrate (A) Fluorogold-labeled spinoparabrachial projection neurons (FG; blue), (B) heat-activated tdTomato (tdT; red) and (C) chloroquine (CQ)-activated Fos (green)-immunoreactive neurons. For multiple labeling, see Supp. Fig. 7. DC: dorsal columns. Scale bars, 100μm.

**Supplementary Figure 6: Retrogradely-labeled projection neurons respond to algogen (heat) and/or pruritogen (chloroquine) stimulation**

Representative sections of lumbar spinal cord from TRAP2-tdTomato mice illustrate Fluorogold-labeled spinoparabrachial projection neurons (FG: blue), heat-activated tdTomato (tdT; red) and/or chloroquine (CQ)-activated Fos-immunoreactive (green) neurons. Arrows point to projection neurons that respond only to CQ (A) or heat (B). Stars point to a projection neuron that responds to both CQ and heat (C). Scale bars, 100μm.

**Supplementary Figure 7: Highly multiplexed *in situ* hybridization reveals widespread subsets of molecularly and functionally diverse projection neurons in the dorsal horn**

**(A)** Representative section of lumbar spinal cord from Retrobead (RB)-injected TRAP2 mouse illustrates merge (A) and individual labeling for DAPI, RB, *Fos* mRNA (heat, 50oC), *tdTomato* mRNA *(tdT;* chloroquine), and genes that mark subsets of projection neurons: *Tacr1, Cck, Nptx2 and Crh.*

