## Supplementary Figures 1-7 for "Pain and itch processing by subpopulations of molecularly diverse spinal and trigeminal projection neurons"

Supplementary Figure 1

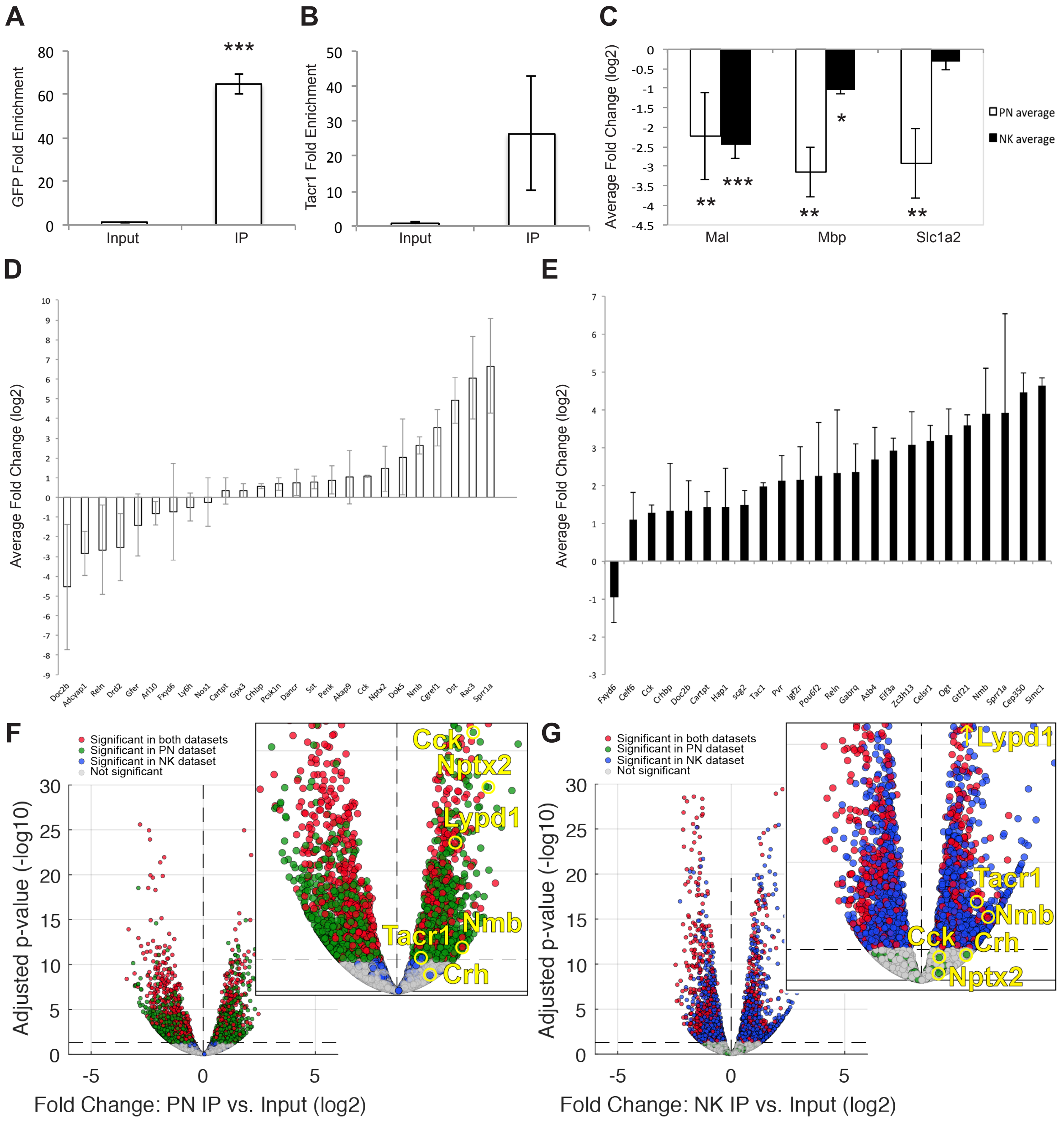

Supplementary Figure 2

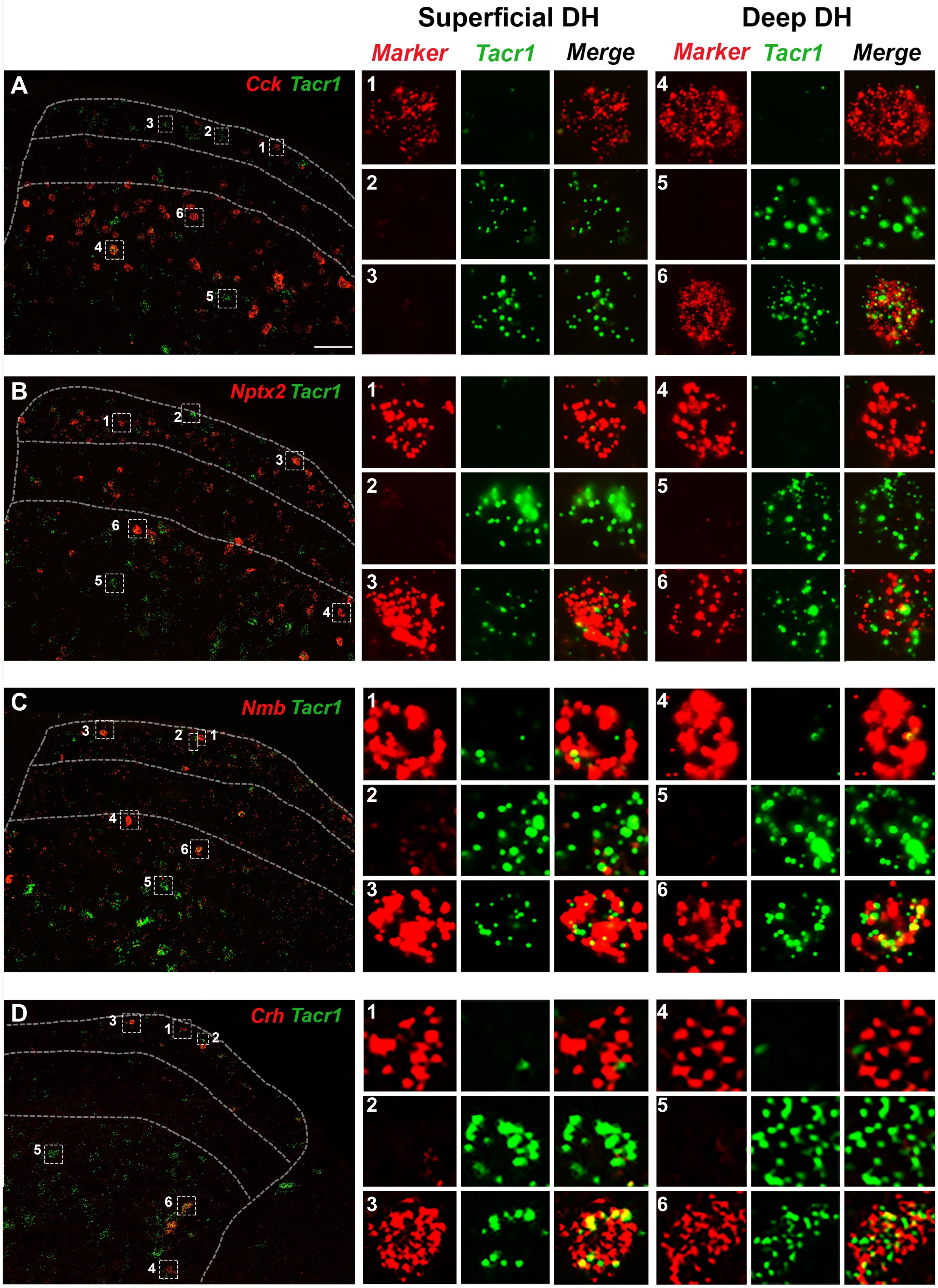

Supplementary Figure 3

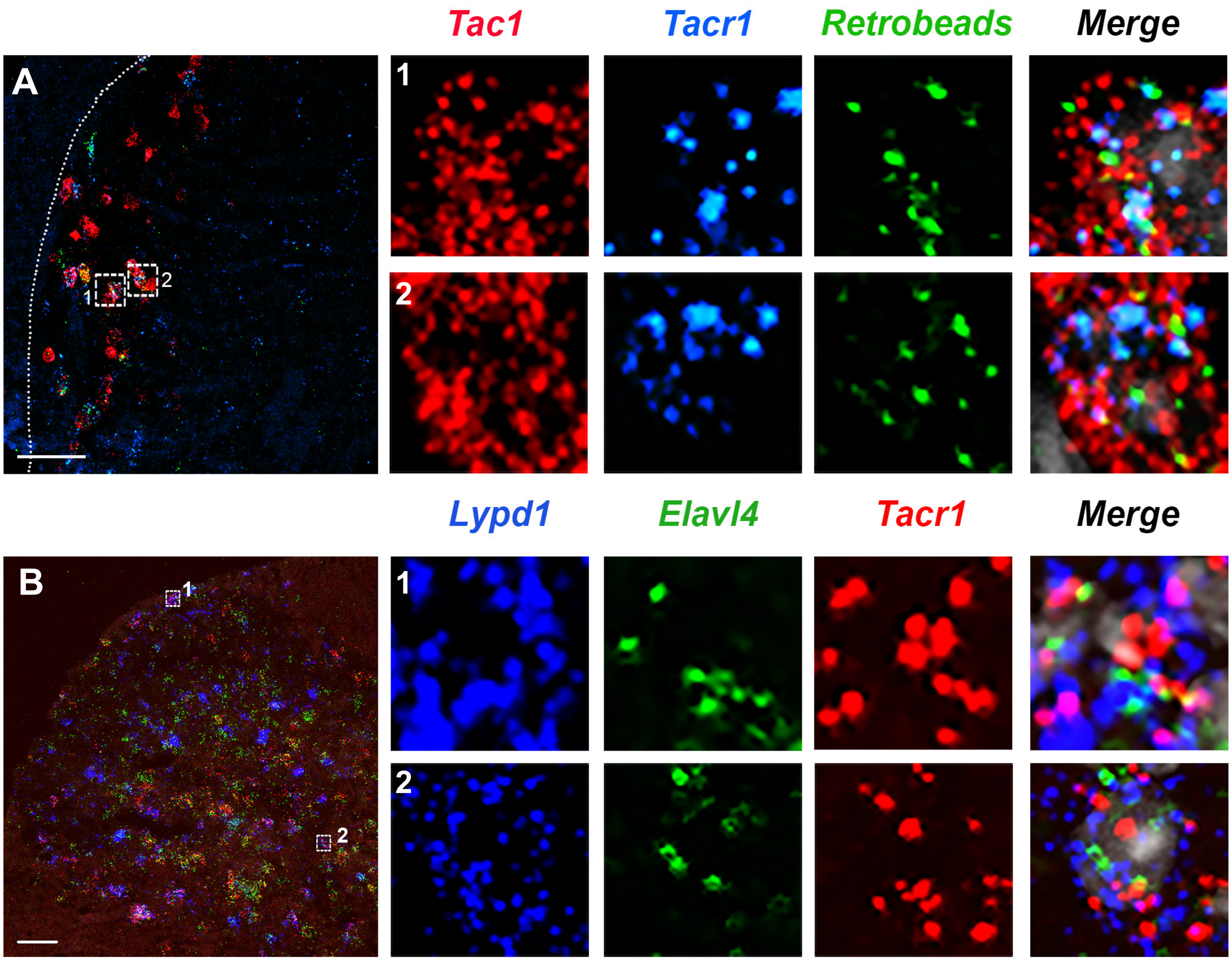

Supplementary Figure 4

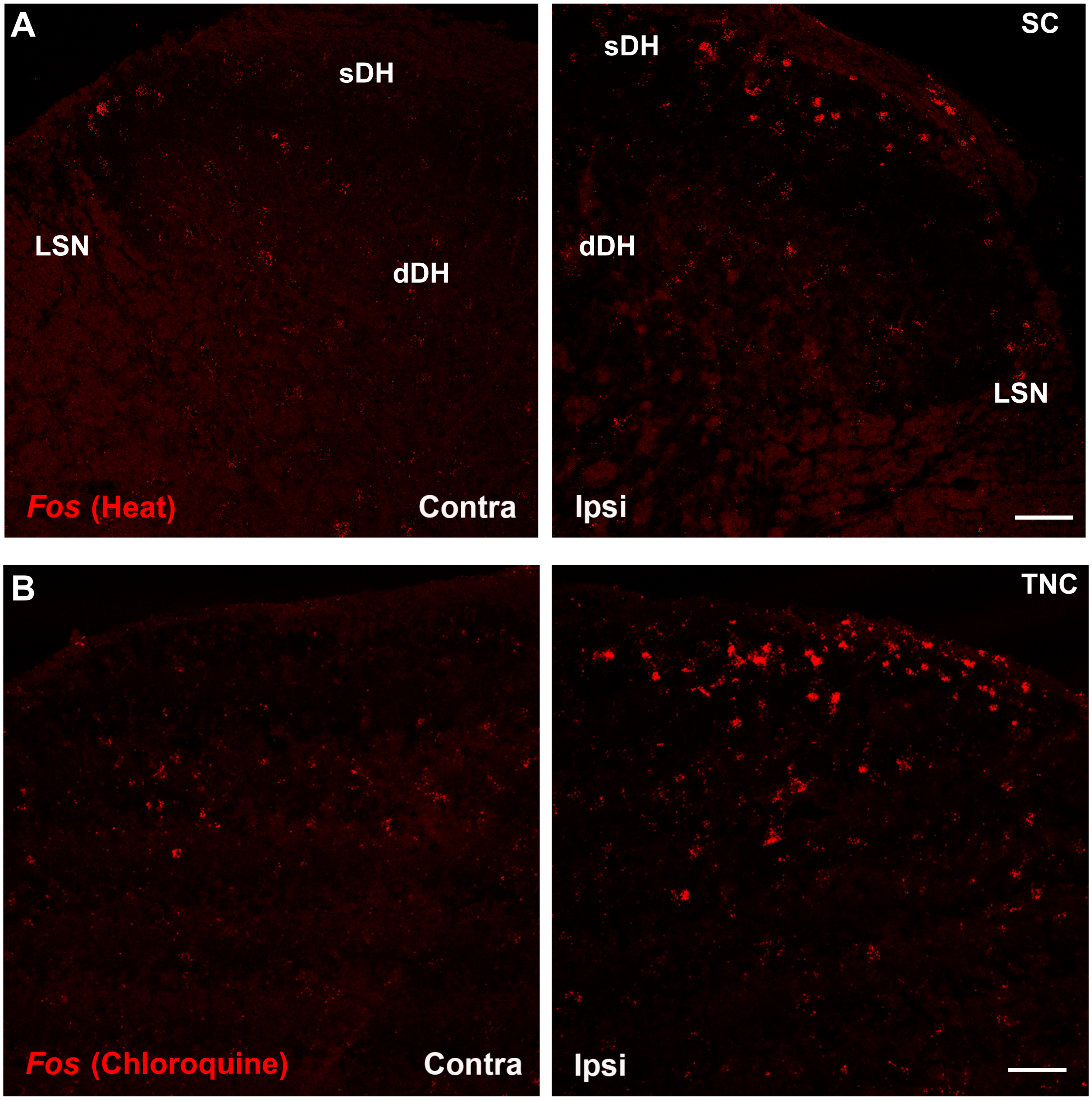

Supplementary Figure 5

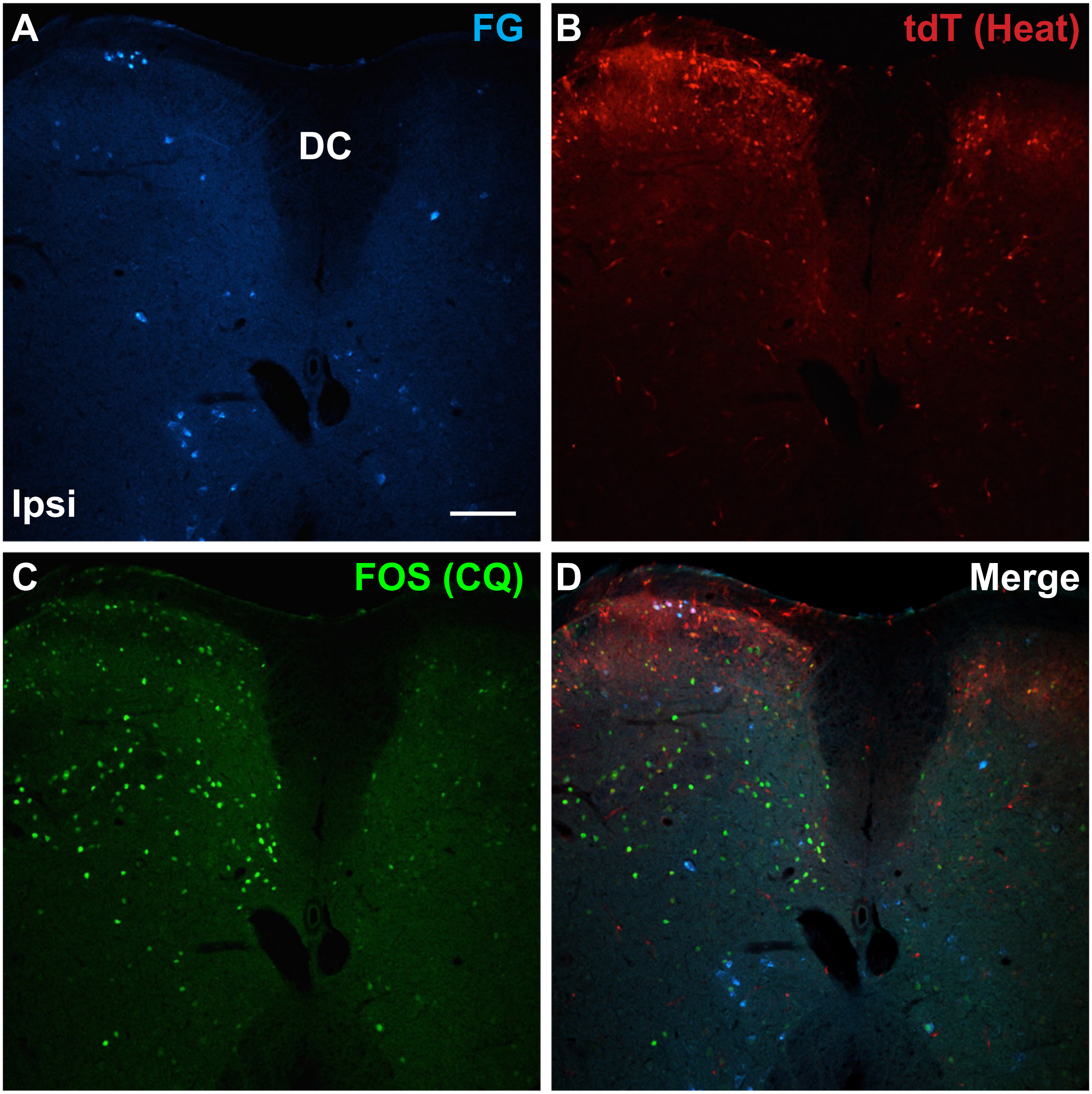

Supplementary Figure 6

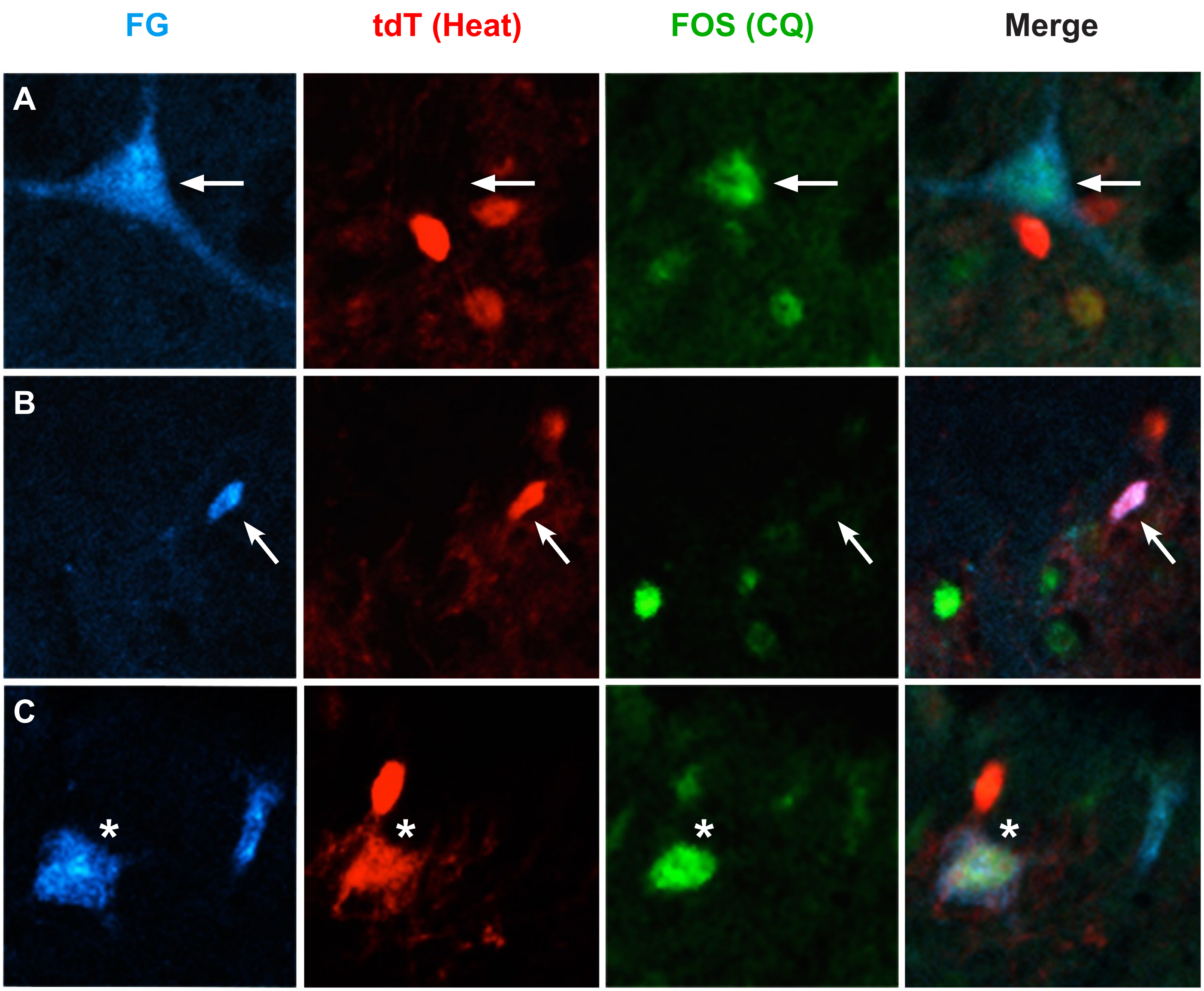

Supplementary Figure 7

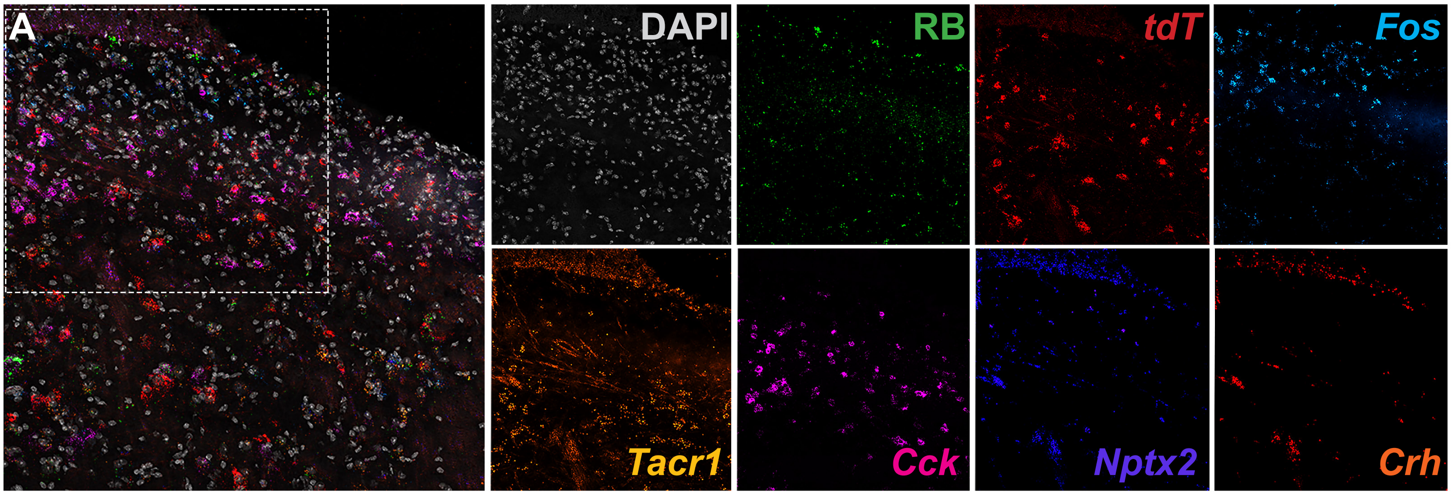
